# Nucleolin is expressed in human fetal brain development and reactivated in human glial brain tumors regulating angiogenesis and vascular metabolism

**DOI:** 10.1101/2020.10.14.337824

**Authors:** Marc Schwab, Ignazio de Trizio, Jau-Ye Shiu, Moheb Ghobrial, Oguzkan Sürücü, Francesco Girolamo, Mariella Errede, Murat Yilmaz, Johannes Haybaeck, Alessandro Moiraghi, Philippe P. Monnier, Sean E. Lawler, Jeffrey Greenfield, Ivan Radovanovic, Karl Frei, Ralph Schlapbach, Viola Vogel, Daniela Virgintino, Katrien De Bock, Thomas Wälchli

## Abstract

Glioblastoma (GBM) is amongst the deadliest human cancers and is characterized by high levels of vascularisation. Angiogenesis is highly dynamic during brain development and almost quiescent in the adult brain, but is reactivated in vascular-dependent CNS pathologies such as brain tumors. Nucleolin (NCL) is a known regulator of cell proliferation and angiogenesis, but its roles on physiological and pathological brain vasculature remain unknown. Here, we studied the expression of Nucleolin in the neurovascular unit (NVU) in human fetal brains and human gliomas *in vivo* as well as its effects on sprouting angiogenesis and endothelial metabolism *in vitro*. Nucleolin is highly expressed in endothelial- and perivascular cells during brain development, downregulated in the adult brain, and upregulated in glioma. Moreover, Nucleolin expression in tumor- and blood vessel cells correlated with glioma malignancy *in vivo*. In culture, siRNA-mediated NCL knockdown reduced human umbilical vein endothelial cell (HUVEC) sprouting angiogenesis, proliferation and filopodia extension, and reduced glucose metabolism. Mechanistically, RNA sequencing of Nucleolin knockdown in HUVECs revealed a putative p53-TIGAR-HK2 regulation of endothelial glycolysis. These findings identify Nucleolin as a neurodevelopmental factor reactivated in glioma that positively regulates sprouting angiogenesis and endothelial metabolism. Our findings have important implications in therapeutic targeting of glioma.

## INTRODUCTION

Glioblastoma (GBM) is a type of glial brain tumor with a putative stem cell origin^1,2^, and represents the most common malignant primary brain tumors^3^. GBMs are amongst the deadliest human cancers, with a less than 15-month median survival and a 5-year survival of only 5%^4–6^. Standard treatment of care involves maximal surgical resection followed by chemotherapy and radiotherapy^6,7^. GBM almost invariably recurs and there is currently no therapy that prolongs survival after tumor recurrence. In order to achieve significant improvements in GBM therapy and clinical outcomes, a better understanding of the cellular and molecular mechanisms that drive GBM growth, propagation, and therapeutic resistance is required.

A typical feature of GBMs is their high grade of vascularization established by angiogenesis, the growth of new blood vessels from pre-existing ones^8–10^. GBM growth is highly dependent on angiogenesis and mutual interaction among the cellular components of the neurovascular unit (NVU)/perivascular niche (PVN) including endothelial- and perivascular cells such as pericytes, astrocytes, neurons, macrophages, microglia, and neuronal stem cells^11^. Accordingly, therapeutic approaches targeting angiogenesis and the NVU in GBM have been proposed^4,12–14^. However, despite promising preclinical data, anti-angiogenic agents have failed to show a survival benefit in randomized controlled trials in GBM patients^4,12^. This is mainly due to our limited knowledge about the cellular and molecular mechanisms regulating angiogenesis and the NVU in brain tumors^4,8,9,11,15,16^.

Whereas angiogenesis is highly dynamic during brain development, the brain vasculature is mostly quiescent in the adult brain with only few proliferating endothelial cells ensuring a stable bloodbrain barrier (BBB)^15,17,18^. Interestingly, angiogenesis and the NVU are reactivated in a variety of angiogenesis-dependent central nervous system (CNS) pathologies such as brain tumors, brain vascular malformations, or stroke^8,11,15,18,19^. However, the molecular signaling cascades reactivated in brain tumors remain elusive. For instance, whether there is molecular similarity between developmental- and tumor angiogenesis, and how neurodevelopmental pathways regulate brain tumor (vessel) growth remains poorly defined. Thus, in order to better understand pathological brain tumor vasculature, a molecular understanding of normal vascular brain development is crucial^15,17,19^.

During development, the brain vascular network is established in multiple steps during embryogenesis as well as at the postnatal stage^15,20,21^. After initial formation of the perineural vascular plexus (PNVP) surrounding the CNS via vasculogenesis (= *de novo* formation of blood vessels from angioblasts), the brain is predominantly vascularized by sprouting angiogenesis (= formation of new blood vessels from pre-existing ones)^15,20,22^. During sprouting angiogenesis, specialized endothelial tip cells at the forefront of a growing vascular sprout extend filopodia to guide growing vessels^15,23,24^. Behind the endothelial tip cells, endothelial stalk cells proliferate to further elongate the vascular sprout ^23,25^, and quiescent endothelial phalanx cells line the parental vessel^18^. After fusion of neighboring sprouts, pericytes and perivascular astrocytes are recruited to stabilize the vessel sprout and allow the formation of functional BBB microvessels^11,12,15,16^. Interestingly, endothelial tip cells drive sprouting angiogenesis during both development and in tumors^8,9,23,26,27^. For instance, endothelial tip cells have been shown to exert crucial regulatory effects via interaction with tumor cells within the perivascular niche in tumors outside the CNS^28,29^. However, even though peri-/neurovascular crosstalk and metabolism are crucial features of brain tumors^7–9,11,12,27,30–34^, less is known about how developmental signaling pathways are reactivated in brain tumors to regulate angiogenesis, perivascular crosstalk, and endothelial metabolism^8,12,30,31^.

Nucleolin (NCL) is a multifunctional and widely expressed protein found in various cell compartments of eukaryotic cells (nucleoplasm, nucleolus, cytoplasm and plasma membrane)^35^. It has been described as the most abundant non-ribosomal component of the nucleolus and its main functions are the regulation of ribosome biogenesis and ribosomal RNA (rRNA) synthesis^36–38^. Moreover, it regulates a variety of other fundamental biological processes such as cell cycle, senescence, apoptosis, and angiogenesis^35,36,39^.

Upregulation of Nucleolin in CNS and non-CNS cancer cells is associated with poor prognosis and promotes cell proliferation and rDNA transcription to sustain a high level of protein synthesis^40–43^. Nucleolin was also shown to prevent breast tumor cell apoptosis by promoting BCL2 expression) and reducing p53 translation^44–46^. Nucleolin expression increases with increasing malignancy grade and proliferation rate in both human and mouse glial brain tumors^43,47–49^, indicating its pro-proliferative role in glioma. Furthermore, Nucleolin was found to be present in GBM stem cells^50,51^, and may constitute a histopathological marker for glioma grading and a possible tool for targeted therapy^51^.

Regarding angiogenesis, Nucleolin was shown to be upregulated at the cell surface of endothelial cells in angiogenic vessels in breast tumor xenografts *in vivo*^52^. At the cell surface, Nucleolin serves as a receptor for various ligands such as Endostatin^53^ and Pleiotrophin^54^, with inhibitory and promoting effects on carcinogenesis and angiogenesis. Moreover, Nucleolin was shown to regulate endothelial cell motility and tube formation *in vitro*^55^, and targeting endothelial cell surface Nucleolin induced endothelial apoptosis and vessel normalization in a pancreatic tumor mouse model^56,57^.

However, the role of Nucleolin in angiogenesis of the developing human brain and in human gliomas CNS has not been investigated and the role of Nucleolin in vascular endothelial cells remains poorly understood. Here, using a variety of *in vivo* and *in vitro* assays, we show that Nucleolin is a neurodevelopmental factor regulating sprouting angiogenesis in the developing CNS and is reactivated in glial brain tumors.

## RESULTS

### Nucleolin is a neurodevelopmental protein that is silenced in the healthy adult brain and reactivated in the NVU/PVN of glial brain tumors

During development, the brain is predominantly vascularized by sprouting angiogenesis ^58^. At the embryonic stage, the perineural vascular plexus in the leptomeninges surrounding the brain is formed via vasculogenesis, followed by the radial growth of penetrating microvessels into the CNS parenchyma^59^. To address the question whether Nucleolin constitutes a neurodevelopmental protein that is reactivated in brain tumors, we performed immunofluorescence microscopy of the main NVU/PVN cellular components of human fetal brain, of human normal adult brain, and human glial brain tumors. Nucleolin was highly expressed during fetal forebrain neocortex development, at gestational week 18 (GW18) and GW22, significantly downregulated in the adult brain, and then upregulated in brain tumors, as revealed by immunofluorescence staining against Nucleolin and the nuclear marker TO-PRO-3^60^ (Fig. 1a-c,p). Moreover, Nucleolin expression was present throughout the nucleoplasm during fetal development but was restricted to the nucleolus in the adult human brain (Fig. 1a,b). In GBM, Nucleolin expression was detected across the entire nucleoplasm and appeared similar to the pattern observed during fetal development (Fig. 1a,c). With regard to its cellular expression within the NVU, Nucleolin was expressed in both endothelial- and perivascular cells (Fig. 1d-o). Nucleolin was highly expressed in cells labeled with the endothelial marker cluster of differentiation 31 (CD31) during brain development, where 84% of CD31^+^ endothelial cells showed Nucleolin expression across the entire nucleoplasm (Fig. 1d, q). Nucleolin was significantly downregulated in the adult brain with 16% of the CD31^+^ cells being Nucleolin^+^ endothelial cells (predominant nucleolar expression) (Fig. 1e,q), but was significantly upregulated in GBM endothelial cells, where 67% of the CD31^+^ were Nucleolin^+^ (nucleoplasm and nucleolar expression), similar to its expression in fetal brain (Fig. 1d,f,q). Nucleolin was also highly expressed in CD105^+^ angiogenic endothelial cells during fetal brain development and in GBM, with 75% and 74% CD105^+^/Nucleolin^+^ double-positive endothelial cells, respectively (Fig. 1g,i,r), whereas only 13% CD105+/Nucleolin^+^ endothelial cells could be observed in adult brain slices (Fig. 1h,r), consistent with the reported quiescence of endothelial cells in the adult normal brain^15,59,61^.

**Figure 1.**
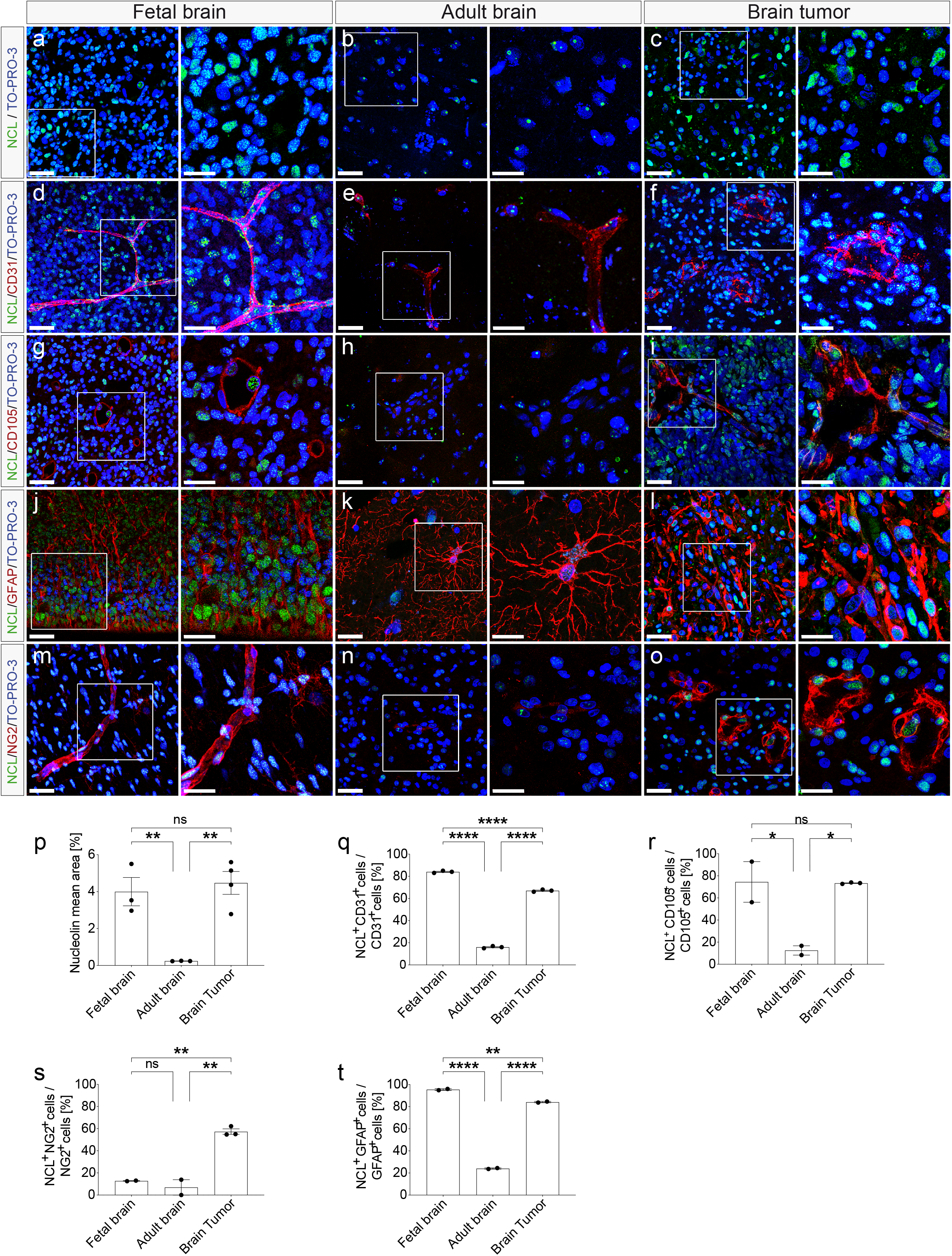
Nucleolin is expressed in endothelial- and perivascular cells during human brain development, is downregulated in the adult brain, and is reactivated in glial brain tumors. Coronal sections (20 μm) of human fetal- (GW 18-22) and adult brain (derived from temporal lobectomies) as well as from human glial brain tumors (GBMs) were stained for Nucleolin (green), the vascular endothelial cell markers CD31 (pan-endothelial marker, red) and CD105 (endoglin, activated endothelial cells, red), the astrocytic marker GFAP (red), the pericyte marker NG2 (red), and TO-PRO-3 nuclear counterstaining (blue). **a,b,c,p** Nucleolin (green) is highly expressed in the nuclei of the developing human fetal brain (**a**) and of human brain tumors (**c**), but shows a significant downregulation in the adult normal/healthy brain (**b**, **p**). **d,e,f,g,h,i,q,r** Nucleolin (green) is highly expressed in CD31^+^ blood vessel endothelial cells (red) in the human fetal (**d,q**)- and pathological brain (**f,q**), but is significantly downregulated in endothelial cell of the quiescent adult brain (**e,q**). Nucleolin shows a high expression in CD105 positive activated endothelial cells (red) in the human fetal brain (**g,r**) and in glioblastoma (**I,r**) but is significantly downregulated in the quiescent adult normal brain (**h,r**), with a very low number of CD105^+^ endothelial cells in the quiescent adult brain (**h,r**). **j,k,l,s** Nucleolin (green) is highly expressed in GFAP^+^ neural precursors cells (red) in the fetal brain (**j**) and in tumoral astrocytes in glioblastoma (**l**) but is downregulated in adult normal brain (**k**, **s**). **m,n,o,t** In human fetal and adult brains, NG2^+^ pericytes (red) partially express low levels of Nucleolin (**m,n**). Nucleolin expression is highly upregulated in human brain tumor NG2^+^ pericytes (**o,t**). (**p-t**) Quantification of Nucleolin expression in all- (**p**), endothelial- (**q,r**), and perivascular cells (**s,t**) in human fetal brain, -adult brain, and -glioblastoma. Nucleolin expression was significantly higher in the developing brain and in brain tumors than in the adult brain, in all cells (**p**), endothelial cells (**q,r**), and astrocytes (**s**). In pericytes, however, Nucleolin expression was significantly higher in brain tumors as compared to brain development and adult brain (**t**). Scale bars: 25 μm in a-o, **left panels**; and 15 μm in a-o, **right panels**. \**P* < 0.05, \*\**P* < 0.01, \*\*\*\**P* < 0.0001.

Within the NVU, blood vessel endothelial cells are in contact with perivascular supportive cells such as pericytes, astrocytes, and neuronal stem cells^15,59,62–65^. Therefore we assessed the expression of Nucleolin in pericytes and perivascular astrocytes. Glial fibrillary acidic protein (GFAP)^+^ astrocytes formed typical patterns by contacting endothelial cells and showed very strong Nucleolin expression during fetal brain development with 95% of GFAP^+^/Nucleolin^+^ astrocytes (Fig. 1j,s). In contrast, in the adult brain, Nucleolin was significantly downregulated in GFAP^+^ astrocytes with only 24% GFAP^+^/Nucleolin^+^ (no restriction to nucleolus observed), but showed a significant upregulation in glioblastoma with 84% of GFAP^+^/Nucleolin^+^ astrocytes (Fig. 1j,k,s). Interestingly, neuron-glial antigen 2 (NG2)^+^ pericytes showed low Nucleolin expression in fetal brain development with only 13% NG2^+^/Nucleolin^+^ pericytes as well as in the adult brain with 7% NG2^+^/Nucleolin^+^ pericytes (Fig. 1m,n,t). However, Nucleolin was significantly upregulated in human glioblastoma with 57% NG2^+^/Nucleolin^+^ pericytes (Fig. 1o,t).

Taken together, these data reveal that Nucleolin is highly expressed in endothelial- and certain NVU cells (astrocytes > pericytes) during fetal brain development, is subsequently downregulated in the adult brain, and is reactivated in GBM. This supports Nucleolin as a neurodevelopmental protein that is reactivated in human glial brain tumors after downregulation in the quiescent adult NVU.

### Nucleolin expression within the NVU correlates with glial brain tumor malignancy and progression

Glial brain tumors are characterized by tumor progression from low- to high grade tumors, described by World Health Organisation (WHO) grades I-IV, whereas gliomas grade I and II are low-grade-, and grade III and IV (= glioblastoma) are malignant (= high-grade) brain tumors^66,67^. In order to address the expression of Nucleolin in glial brain tumor progression, we referred to tissue microarrays (TMAs) of human glioma stained immunohistochemically for Nucleolin and the nuclear marker Mayer’s hemalum (Fig. 2a-d). Nucleolin expression was markedly upregulated in human glial brain tumors as compared to the adult normal brain (Fig. 2a-d, e). Moreover, Nucleolin expression was significantly increased during glial tumor progression, ranging from 32% of Nucleolin^+^ cells in WHO grade I glioma to 57% in glioma grade IV (= glioblastoma) (Fig. 2e). Nucleolin showed a significant upregulation from low grade (WHO grade I and II) to high grade glioma (WHO grade III) as well as a significant increase from WHO grade III to WHO grade IV glioma (= glioblastoma) (Fig. 2a-d, f). In recurrent glioblastoma, Nucleolin expression showed a slight but significant decrease in expression as compared to primary glioblastoma (Fig. 2e). Nucleolin expression correlated well with the established proliferation marker Ki-67^68^ in all glioma grades (Fig. 2g).

**Figure 2.**
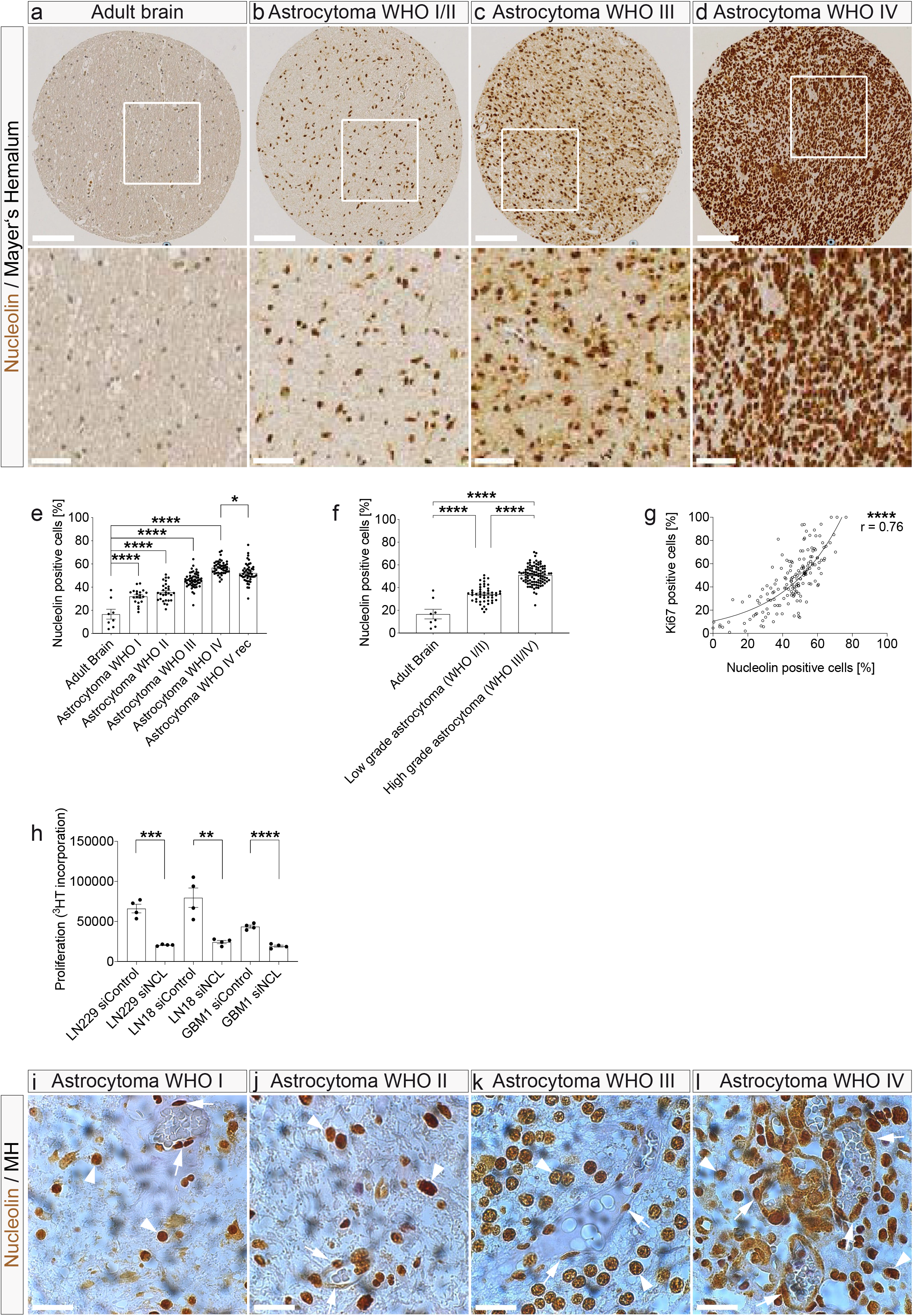
Expression of Nucleolin increases during astrocytic tumor progression and Nucleolin is expressed in blood vessels in low and high grade gliomas. **a-f** Nucleolin (brown) expression increases during tumor progression of human astrocytomas from WHO grade I (**a**), II (**b**), III (**c**) to IV (= glioblastoma, **d**). In low-grade astrocytomas (WHO grades I and II), Nucleolin expression (as revealed by the percentage of Nucleolin^+^ cells) is significantly higher than in the normal brain parenchyma (**e,f**), but significantly lower as compared to high-grade astrocytomas (WHO grades III and IV) (**e,f**). Nucleolin expression slightly decreases again in recurrent WHO grade IV tumors (**e**). **g** Nucleolin expression positively correlates with the expression of the proliferation marker Ki-67. **h** Glioblastoma cell lines (LN229, LN18 and GBM-1) proliferation was significantly decreased upon Nucleolin knock down, as revealed by the reduced [3H-methyl] thymidine [(3H]TdR) incorporation. **j-m** Nucleolin expression in tumor blood vessels. Note the increasing expression of Nucleolin in the blood vessel wall (arrows) as well as in perivascular cells (arrowheads) in astrocytomas of higher grades. \*\**P* < 0.01, \*\*\*\**P* < 0.0001. Scale bars represent: 100 μm (**a-d**, upper panel), 50 μm (**a-d**, lower panel, and **g-j**).

Based on Nucleolin’s expression within perivascular cells of the developmental- and tumoral NVU, we first aimed to address its effects on tumoral cell proliferation. To determine whether Nucleolin promotes GBM cell proliferation, Nucleolin was knocked down in the human glioblastoma cell lines LN-229^69^ and LN-18^70^ as well as in freshly isolated primary human glioblastoma cells (GBM-1) using siRNA (data not shown). Cell proliferation of LN-229, LN-18, and GBM-1 was inhibited by siRNA-mediated knockdown of Nucleolin (Fig. 2h) compared with scrambled controls, in agreement with the previously reported strong pro-proliferative effect of Nucleolin in human glioblastoma cells^43,71^.

Next, we examined the immunohistochemical expression of Nucleolin in tumor blood vessels. In agreement with our immunofluorescent data in GBMs (see Fig. 1f,y), Nucleolin was indeed present in the wall of tumor blood vessels, showing an increased expression during glial tumor progression (WHO I – IV, Fig. 2i-l), further suggesting a regulatory effect on human glial brain tumor angiogenesis.

Taken together, these data indicate that Nucleolin expression is reactivated in tumor- and tumor-associated endothelial cells within the NVU during human astrocytic tumor progression.

### Nucleolin is expressed within the NVU and in sprouting endothelial tip cells, and promotes the number of tip cell filopodia during brain development *in vivo*

Nucleolin has been shown to affect tumor growth and angiogenesis^43,52,55,71,72^, but the underlying cellular and molecular mechanisms of its roles in the NVU/PVN and on sprouting angiogenesis during brain development remain unclear.

To assess whether Nucleolin affects sprouting angiogenesis and endothelial tip cells during brain development, we addressed Nucleolin expression in the vicinity of sprouting blood vessels in the human fetal brain. CD105-labeled endothelial tip cells with their typical, finger-like protruding filopodia could be recognized in GW 18 and 22 human fetal brain forebrain (Fig. 3a-d). Nucleolin was expressed in CD105^+^ endothelial tip-, stalk-, and phalanx cells (Fig. 3a-d) as well as in perivascular cells surrounding sprouting capillary endothelial tip cells (filopodia) (Fig. 3a-d). We observed Nucleolin in nuclei of CD105^+^ endothelial tip-, stalk-, and phalanx cells but not on the (endothelial- and perivascular) cell surface and not in filopodia protrusions (Fig. 3a-d).

**Figure 3.**
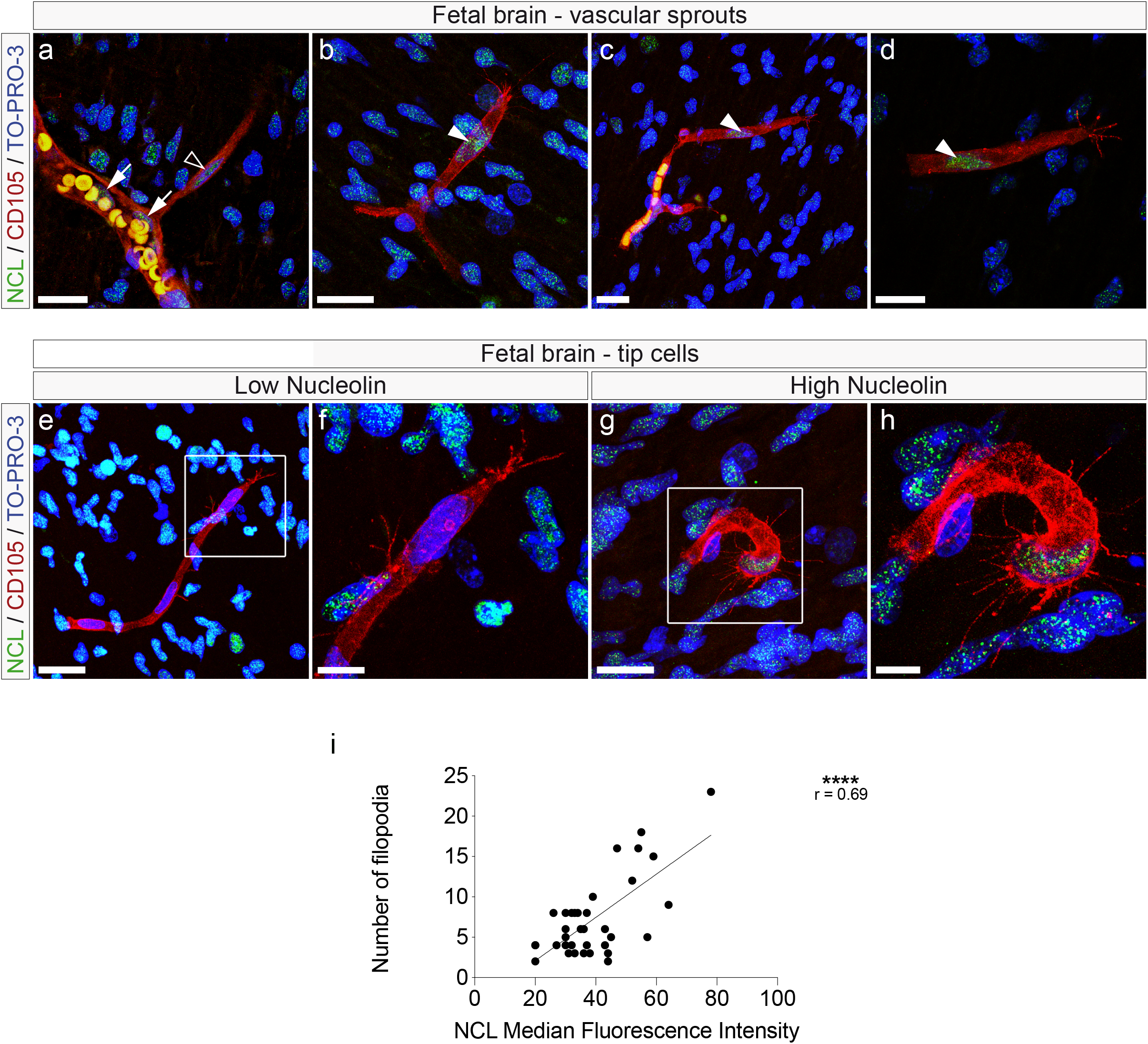
Nucleolin is expressed in endothelial tip-, stalk-, and phalanx cells of vascular sprouts and Nucleolin expression correlates with the number of endothelial tip cell filopodia in the human fetal brain *in vivo*. 20 μm sections of human fetal (GW 18-22) brains were stained for Nucleolin (green), the vascular endothelial cell marker CD105 (red) and TO-PRO-3 nuclear counterstaining (blue). **a-d** Blood vessel sprouts (red) in the human fetal brain grow in CNS tissue with vascular- and parenchymal expression of Nucleolin, respectively. Nucleolin (green) is expressed in endothelial tip (filled arrowheads in **b**, **c,d**)- stalk (empty arrowhead in **a**)- and phalanx cells (plain arrows in **a**) in growing vessels in the human fetal (GW18-22) cortex. **e-h** Vascular sprouts with CD105-labeled endothelial tip cells (red) with low- (e,f) and high (g,h) Nucleolin (green) expression in the human fetal cortex. Numerous filopodial protrusions emerged from the endothelial tip cell body with high Nucleolin expression (**g,h**) as compared to only few filopodial protrusions in an endothelial tip cell with low Nucleolin expression (**e,f**). (**i**) The number of endothelial tip cell filopodia strongly correlated with the intensity of Nucleolin expression. \*\*\*\**P* < 0.0001. Scale bars: 20 μm in a-d; 25 μm in e and g; and 10 μm in f and h.

We examined if Nucleolin expression affected the number of endothelial tip cell filopodia, and observed a lower number of filopodia in endothelial tip cells with low Nucleolin expression (Fig. 3e,f) and a higher number of filopodia in endothelial tip cells with high Nucleolin expression (Fig. 3g,h). Accordingly, we quantified the number of filopodia in Nucleolin^+^/CD105^+^ endothelial tip cells and assessed Nucleolin expression for each tip cell. Indeed, the number of filopodia positively correlated with Nucleolin expression in the tip cells, as revealed by median fluorescent intensity (Fig. 3i).

These results strongly suggest that Nucleolin positively regulates the number of endothelial tip cell filopodia in the human fetal brain.

### Nucleolin promotes HUVEC sprouting angiogenesis *in vitro*

Based on these expression studies suggesting a role for Nucleolin in sprouting angiogenesis and endothelial tip cell filopodia *in vivo*, we next investigated the functional role of Nucleolin in human angiogenic endothelial cell sprouting *in vitro*. We used siRNA, to knock-down Nucleolin in human umbilical vein endothelial cells (HUVECs), and stained them with phalloidin to reveal the F-actin cytoskeleton (Fig. 4a-d). In Nucleolin siRNA-treated HUVECs (HUVEC^Nucleolin KD^), Nucleolin expression was decreased and confined to the nucleolus when compared to the siRNA control-treated HUVECs (HUVEC^Control KD^, Fig. 4a-d). Accordingly, qRT-PCR and Western blot analysis revealed significant knocking-down of Nucleolin at both the mRNA- and the protein levels in HUVECs (Fig. 4e-g).

**Figure 4.**
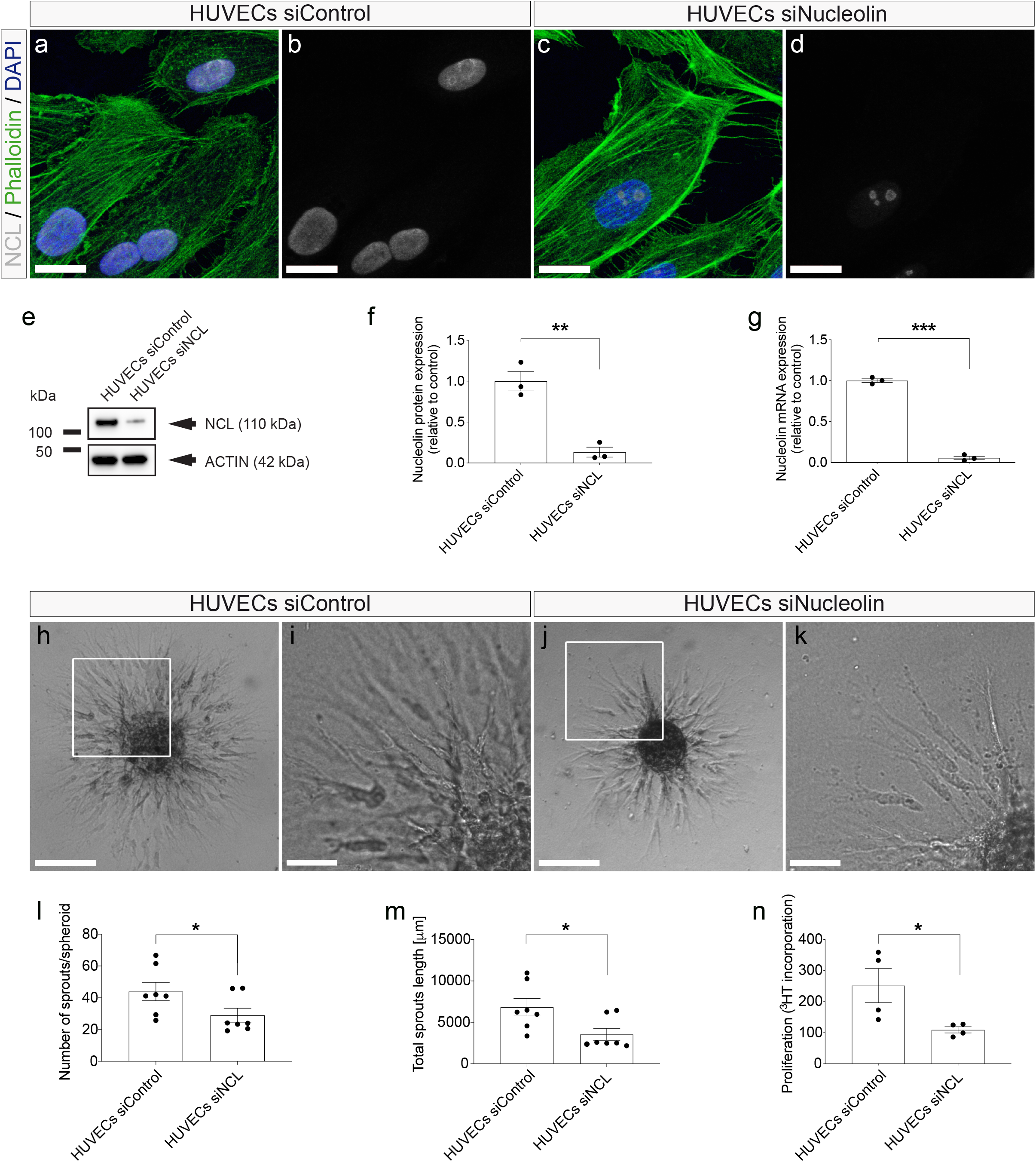
Nucleolin promotes vascular endothelial cell sprouting and proliferation *in vitro*. **a-d** HUVECs were stained for Nucleolin (green), F-actin (stained with Phalloidin, red), and the general nuclear marker DAPI (blue). Nucleolin expression was decreased and restricted to nucleoli upon siRNA-mediated knockdown in HUVECs (**c,d**). No inhibition of Nucleolin expression and no restriction to nucleoli in HUVECs could be seen in the HUVECs treated with a control siRNA (**a,b**). **e** Western blot showing Nucleolin downregulation in HUVECs transfected with siRNA against Nucleolin (siNCL). No Nucleolin downregulation was observed in HUVECs transfected with the control siRNA (siControl). **f** Quantification of Western blot revealing a significant downregulation of 80% Nucleolin protein expression by siRNA-targeted Nucleolin knock down as compared to control cells. **g** Quantitative RT-PCR revealing a significant downregulation of more than 90% of Nucleolin mRNA expression by siRNA-targeted Nucleolin knock down. **h-k** HUVEC sprout formation using hanging drops composed of a collagen type I-methylcellulose matrix containing VEGF-A, basic FGF (bFGF), and EGF was massively decreased upon siRNA-mediated Nucleolin knockdown (**j,k**) as compared to the control group (**h,i**). The boxed areas in **h,j** are enlarged in **i,k** respectively. **l**, **m** Nucleolin downregulation reduces the sprout formation of HUVECs as compared to the control groups. HUVEC sprout formation (number of sprouts per spheroid) and total sprout length were significantly reduced upon Nucleolin knock-down as compared to the control group (**l,m**). finished figure legend Figure 5 from Wälchli et al., PNAS, 2013 paper **n** HUVEC proliferation was significantly decreased upon Nucleolin knock down, as revealed by reduction of radioactively labeled Thymidine incorporation. \**P* < 0.05, \*\**P* < 0.01, \*\*\**P* < 0.001, \*\*\*\**P* < 0.0001. Scale bars: 20 μm in a-d; 150 μm in h and j; and 50 μm in i-k.

To test the effects of Nucleolin on sprouting angiogenesis *in vitro*, we referred to an *in vitro* spheroid angiogenesis assay^73^. HUVECs in the sprouting spheroid assay using hanging drops composed of collagen type I-methylcellulose matrix containing VEGF-A, FGF and EGF grew vessel-like sprouts composed of multiple branches in the control group (Fig. 4h,i). In contrast, siRNA-mediated knock-down of Nucleolin markedly suppressed the number of vessel sprouts per spheroid as well as the length of the sprouts as compared to the control group (Fig. 4j,k,l,m,n,o), suggesting a promoting effect on sprouting angiogenesis.

Given the important role of Nucleolin in cell proliferation^35,37,38^, we next assessed whether Nucleolin affects endothelial cell proliferation in a 3HT proliferation assay. Indeed, HUVEC proliferation was significantly reduced upon siRNA-mediated Nucleolin knock-down (Fig. 4o), indicating a positive regulatory role for Nucleolin on HUVEC cell proliferation, reminiscent of stalk cell behavior *in vivo*^15,23,59^.

Taken together, these results suggest that endothelial Nucleolin is a positive regulator of sprouting angiogenesis, endothelial proliferation-, and filopodia formation in the brain.

### Nucleolin regulates HUVEC lamellipodia and filopodia formation and actin cytoskeleton orientation

Lamellipodia and filopodia are composed of actin and myosin fibers, and are essential components of *in vivo* sprouting angiogenesis^24^. Therefore, to further assess the effects of Nucleolin on the HUVEC angiogenesis *in vitro*, we addressed cell shape and morphology as well as actin orientation after Nucleolin knockdown (Fig. 5a-i). HUVEC^Nucleolin KD^ spread less and were not as elongated as control cells as quantified by their increased cell circularity and decreased cell area (Fig. 5g,h). FActin fibers were more randomly organized in the HUVEC^Nucleolin KD^ (Fig. 5l-p). Accordingly, the distribution of actin orientation showed a classical peak in controls, whereas in the HUVEC^Nucleolin KD^ the actin orientation was more randomly distributed (Fig. 5s), indicating the HUVEC^Nucleolin KD^ had poorly orientated stress fibers as opposed to well-aligned stress fibers of control cells.

**Figure 5.**
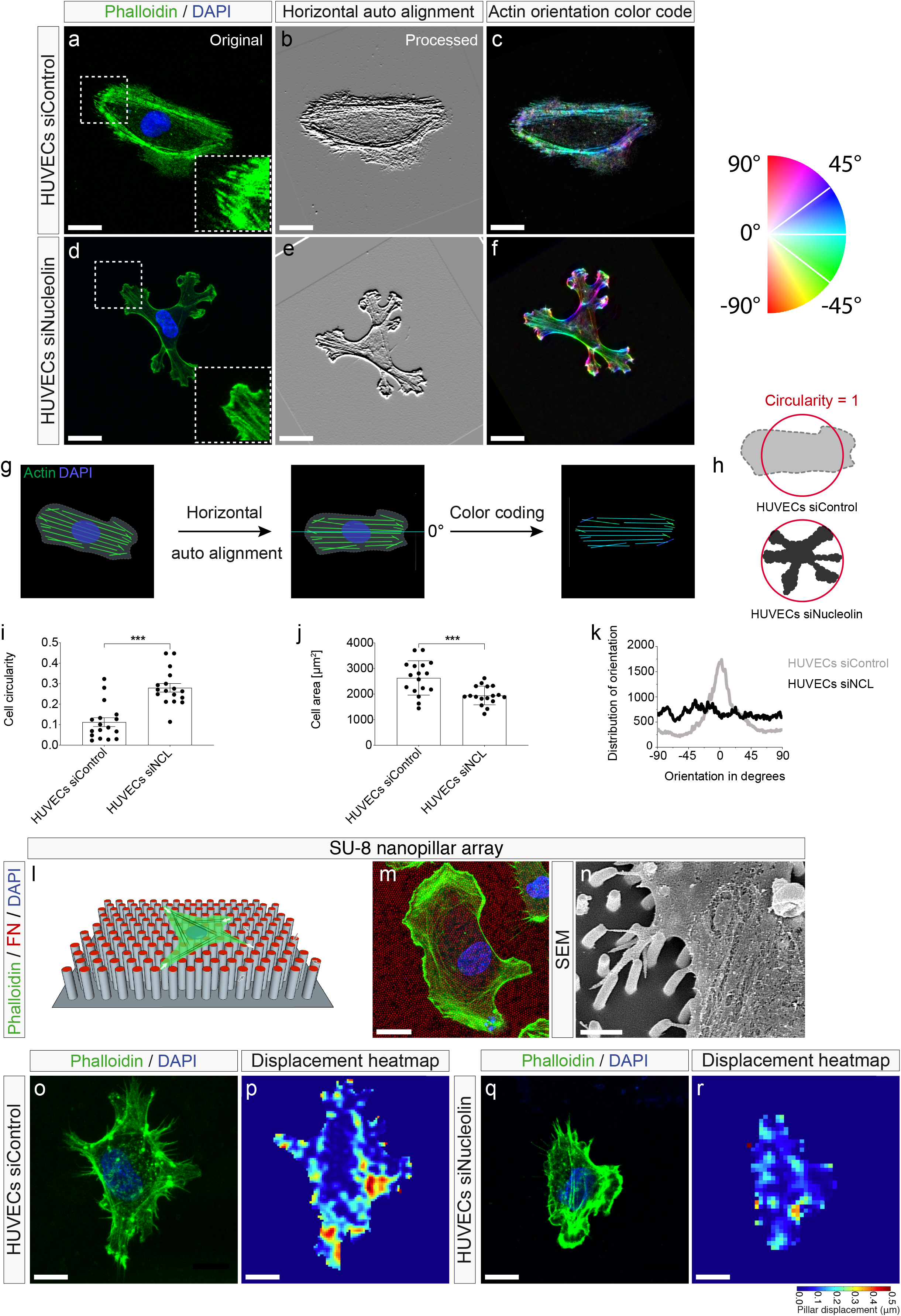
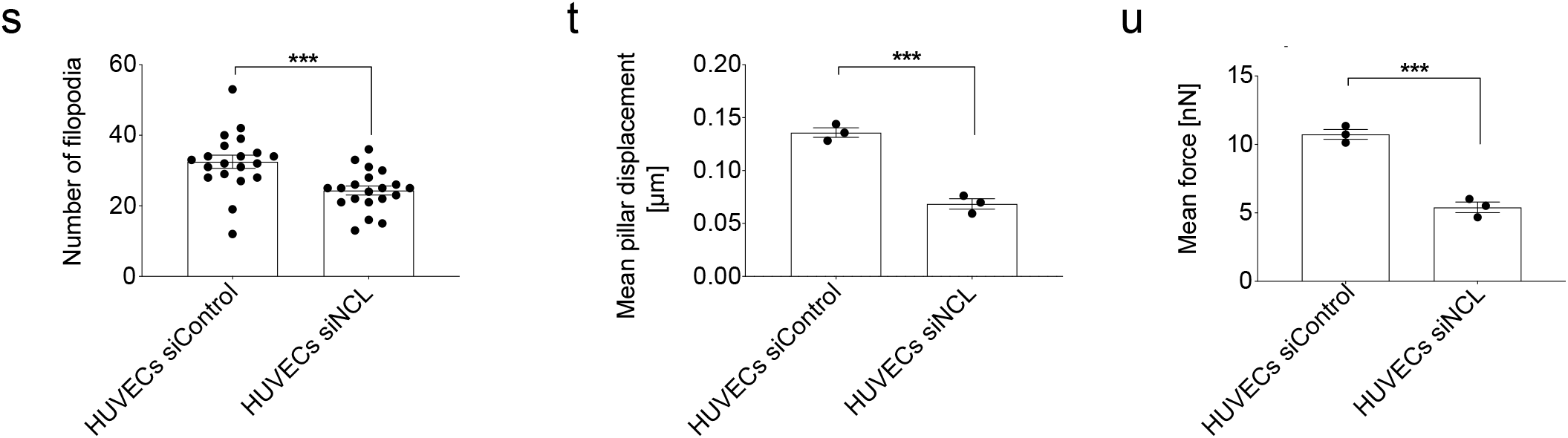
Nucleolin regulates HUVEC lamellipodia and filopodia and affects actin cytoskeleton orientation. **a-f** HUVEC treated with control- or Nucleolin siRNA were left to spread on fibronectin-coated glass substrate and stained for F-actin (green) and DAPI (blue) (**a,d**). Pictures were processed using OrientationJ (plugin of ImageJ) to automatically orientate a maximum of the actin fibers of the cell to a 0° line. Each fiber was then colored based on the color wheel. **g** Scheme illustrating actin fiber orientation characterization using the OrientationJ. **h** Schematic illustration of circularity index. **i,j** Nucleolin knockdown decreased HUVEC cell spreading, as measured by cell circularity and cell area measurements. Nucleolin knock down HUVECs tend to have a less elongated shape as revealed by their circularity coefficient (**i**). HUVEC spreading was decreased upon Nucleolin knockdown as revealed by a lower cell area (**j**). Phalloidin actin fibers (green) were more randomly organized in the Nucleolin-KD HUVECs (**e,f**) as compared to the control (**b,c**). The distribution of actin orientation shows clear classical peak close to 0 degree in the control HUVECs (grey curve). In siNucleolin HUVECs, the classical peak of actin orientation was lost and HUVEC actin orientation was more randomly distributed (black curve) (**k**). **l-n** Scheme illustrating a spread endothelial cell (green) on a SU-8 nanopilllar array (grey) coated with fibronectin (red) (**l**). F-actin (green) and DAPI (blue) stained HUVEC on a fibronectin-coated nanopillar substrate (red) (**m**). Scanning electron microscopy (SEM) image of HUVEC filopodia attaching to nanopillars (**n**). Note the nanopillar-deflection caused by retracting HUVEC filopodia (arrowheads), allowing to optically measure the displacement of the nanopillar and the induced corresponding traction forces. **o-u** HUVECs treated with siRNA (for Nucleolin, and control) were placed on nanopillar substrate, and were stained for F-actin (green) and the general nuclear marker DAPI (blue) (**o**, **q**). The number of filopodia per cell was decreased significantly in siRNA-mediated Nucleolin knockdown in comparison with control siRNA-treated HUVECs (**s**). Explorative movements of HUVECs (and its lamellipodia- and filopodia extensions) were reduced upon Nucleolin knockdown as evidenced by displacement heatmaps (**p,r**). Nucleolin downregulation decreased mean nanopillar displacement (**t**) and mean filopodia force (**u**) accordingly Scale bars: 100 μm in a; 20 μm in d; 2 μm in e; 10 μm in f,g, and k); and 5 μm in h-j, and l-n. \*\*\**P* < 0.001.

To assess the effects of Nucleolin on HUVECs and their filopodia, we cultured these cells on a substrate consisting of nanopillar arrays^74,75^ (Fig. 5j-l) allowing assessment of filopodia dynamics and traction/pulling forces exerted by the spreading of HUVECs on the substrate (Figure. 5k,l). In HUVEC^Nucleolin KD^, the number of filopodial extensions per HUVEC was significantly reduced as compared to HUVEC^Control KD^ (Fig. 5m,o,q).

Next, we examined the movement of Nucleolin-siRNA treated HUVECs on the nanopillar surface. In HUVEC^Nucleolin KD^, the mean displacements of the nanopillars was significantly decreased as compared to the control group (0.068 μm and 0.136 μm, respectively, Fig. 5n,p,r). HUVEC^Nucleolin KD^ exerted significantly reduced average traction forces of 5.4 nN when compared to the control HUVECs (10.7 nN Fig. 5s). These results indicate a pro-adhesive / pro-migratory / pro-explorative effect of Nucleolin on endothelial cells and their filopodial protrusions.

These data indicate that Nucleolin is important for actin orientation and polarization which is required for endothelial cell lamellipodia- and filopodia formation, structures that are crucial for migration, proliferation, and sprouting of vascular endothelial cells *in vivo*.

### Nucleolin regulates endothelial glucose-, but not fatty acid metabolism

Endothelial metabolism has been shown to be a crucial regulator of sprouting angiogenesis, endothelial tip cell formation, and endothelial lamellipodia- and filopodia-dynamics^76–79^. Moreover, endothelial cell glycolysis regulates the rearrangement of endothelial cells by promoting filopodia formation and by reducing intercellular adhesion ^79^. Based on the observed regulatory effects of Nucleolin on sprouting angiogenesis, endothelial tip cell (filopodia) and the actin cytoskeleton, we investigated whether Nucleolin affected endothelial glucose and fatty acid metabolism^78^.

Using a glycolytic flux assay^80^, siRNA-mediated knock-down of Nucleolin resulted in a significant reduction of glycolysis as compared to the control HUVECs (Fig. 6a). Similarly, HUVEC^Nucleolin KD^ showed reduced glucose uptake and decreased lactate production as compared to the HUVEC^Control KD^ (Fig. 6b,c), indicating a positive regulatory effect of Nucleolin on HUVEC glucose metabolism.

**Figure 6:**
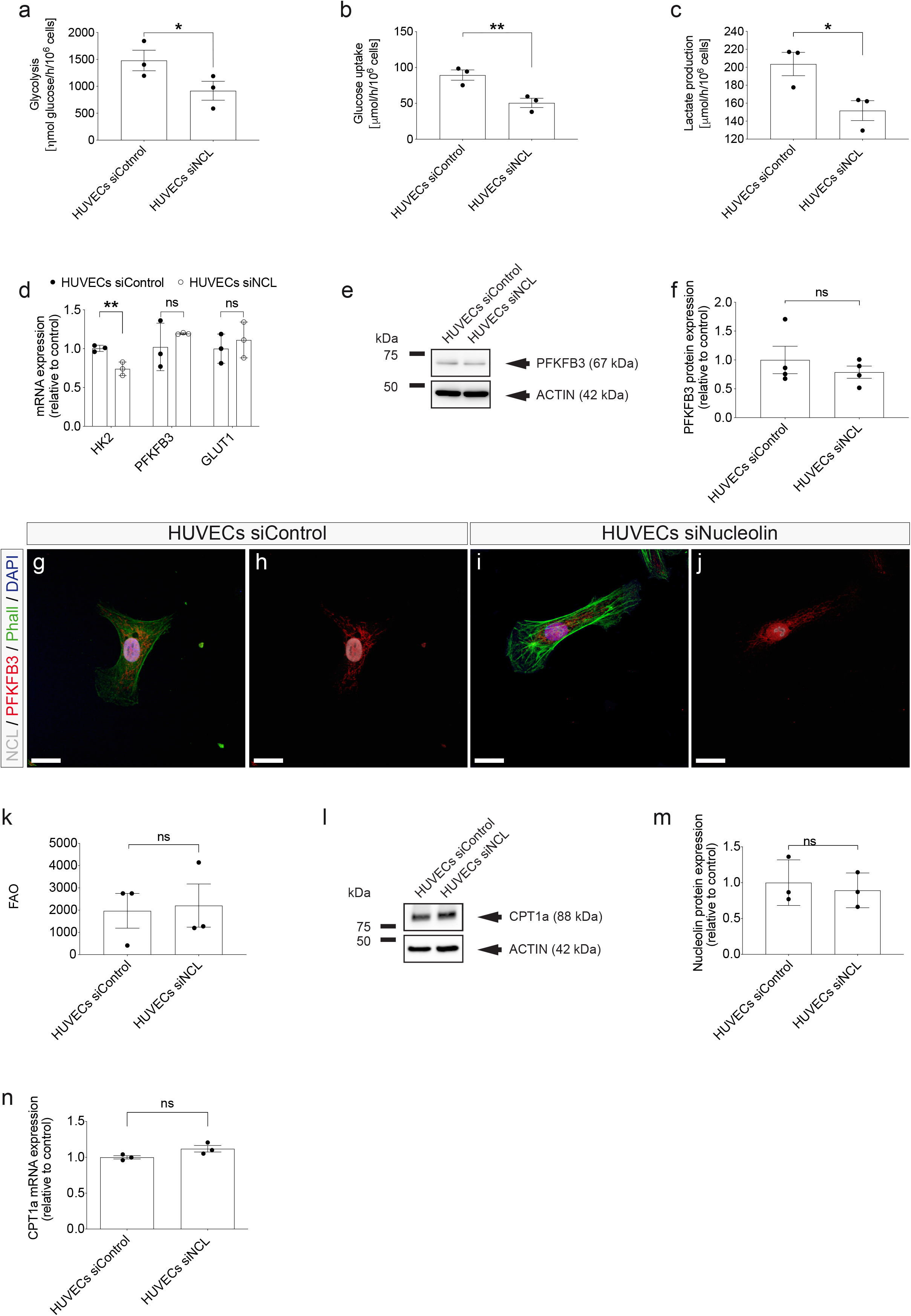
Interaction of Nucleolin signaling with metabolic pathways in human endothelial cells. **a**-**c** Metabolic assays of HUVECs, upon Nucleolin downregulation with small interfering RNA targeting Nucleolin. Nucleolin knockdown decreased the glycolytic flux (**a**), glucose uptake (b) and lactate production (**c**) in HUVECs as compared to the tested controls. **d** Quantitative RT-PCR revealing a significant downregulation of about 30% of Hexokinase-2 (HK2) mRNA expression by siRNA-targeted Nucleolin knock down. 6-phosphofructo-2-kinase/fructose-2,6-biphosphatase 3 (PFKFB3) expression showed slight but no significant increase. **e**-**f** Western blot using antibodies against HK2 and PFKFB3 revealed a significant downregulation of HK2, while PFKFB3 expression remained unchanged by Nucleolin knock down. **g-n** HUVECs were stained for Nucleolin (green), PFKFB3 or HK2 (red), and the general nuclear marker DAPI (blue). HK2 expression was decreased in HUVECs^Nucleolin KD^ (**m,n**) as compared to HUVECs^Control KD^ (**k**, **l**). No difference in PFKFB3 expression could be seen between Nucleolin knock down HUVECs (**i**, **j**) and the HUVECs treated with a control siRNA (**g,h**). **o** Nucleolin knock down in HUVECs did not affect fatty acid oxidation. **p** Quantitative RT-PCR showed no significant regulation of carnitine palmitoyltransferase 1A (CPT1a) mRNA expression upon siRNA-targeted Nucleolin knock down. **q**-**r** Western blot using antibodies against the CPT1a showed no difference in CPT1a protein expression between Nucleolin knock down HUVECs and the control condition. \**P* < 0.05, \*\**P* < 0.01.

6-phosphofructo-2-kinase/fructose-2,6-biphosphatase 3 (PFKFB3), and Hexokinase2 (HK2), have been shown to be key regulators of endothelial glucose metabolism^77,78^. To address whether Nucleolin affected the expression patterns of these genes involved in endothelial glucose metabolism in HUVECs, we performed qRT-PCR and Western blots of HUVEC^Nucleolin KD^ and HUVEC^Control KD^. HK2 was significantly downregulated upon Nucleolin knockdown on the mRNA level (Fig. 6d), whereas PFKFB3 showed no significant change on both the mRNA and the protein levels (Fig. 6d-j).

Next, we assessed whether Nucleolin also regulates endothelial fatty acid oxidation (FAO) which was also shown to exert crucial effects endothelial stalk cells proliferation^78,81^, and to be upregulated in quiescent endothelial cells as a protection against oxidative stress^82^. Nucleolin KD in HUVECs did not affect endothelial fatty acid oxidation (Fig. 6k). Carnitine palmitoyltransferase 1A (CPT1a) has been shown to be a key regulator of endothelial fatty acid metabolism^77,78^. As expected, qRT-PCR and Western blot analysis of HUVEC^Nucleolin KD^ showed no significant differences of CPT1a as compared to the HUVEC^Control KD^ group at both the mRNA- and protein-levels (Fig. 6l,m,n).

Taken together, these data reveal that KD of Nucleolin decreases endothelial glucose metabolism without affecting endothelial fatty acid metabolism, indicating a positive regulatory role of Nucleolin on sprouting angiogenesis via promoting endothelial glucose metabolism.

### Bulk RNA sequencing (RNAseq) reveals upregulation of PFKFB3-independent endothelial glycolysis regulators p53 and TIGAR

To address the downstream signaling pathways induced by Nucleolin in human endothelial cells, we performed an unbiased transcriptome analysis by RNAs of HUVEC^Nucleolin KD^ and HUVEC^Control KD^ (Fig. 7a-c). RNAseq revealed 3240 genes significantly differentially regulated genes between HUVECs^siRNA-Nucleolin KD^ and HUVECs^siRNA-control KD^ cells. (Fig. 7a-c). Among the top-regulated genes upon Nucleolin KD figured genes implicated in cell proliferation and cell cycle progression, in agreement with the previously reported role of Nucleolin in these cellular processes^37,38,43^. To that regard, the key cell cycle inhibitor CDKN1A/P21 figured within the top upregulated genes, whereas several regulators of the cell cycle such as the mitotic spindle protein cytoskeleton associated protein 2 like (CKAP2L), DNA topoisomerase 2-alpha (TOP2A), mitosis regulator DEPDC1 and cyclin dependent kinase 1 (CDK1) were the top downregulated genes in HUVECs^Nucleolin KD^ (Fig. 7d). Analysis of the gene expression by the fragments per kilobase of exon model per million reads mapped (FPKM) values, confirmed the Nucleolin downregulation mediated by siRNA (Fig. 7e). Based on the observed regulatory effect of Nucleolin on sprouting angiogenesis, we analyzed the expression of main regulators of key angiogenic pathways such as the VEGF-A/VEGFR2/3-Dll4-Jagged-Notch-, and the Hippo-YAP-TAZ-pathways (Fig. 7f-h). Indeed, HUVEC^Nucleolin KD^ expressed higher levels of the Notch downstream effectors HES1 and HES2, as well as of the NOTCH ligand Jagged 1 (JAG1) (Fig. 7f). On the other hand, Nucleolin knock-down induced a downregulation of the expression of Notch receptor 4 (NOTCH4) and of the Notch ligand DLL4 (Fig. 7f). Moreover, Nucleolin knock-down downregulated VEGFA but upregulated VEGFR2 (Fig. 7g). Within Hippo-YAP-TAZ pathway, YAP1 expression and downstream effectors CTGF and CYR61 were significantly downregulated with Nucleolin knockdown (Fig. 7h). The upregulation of the anti-angiogenic Dll4-Notch as well the downregulation of the pro-angiogenic Hippo-YAP-TAZ pathway upon Nucleolin-KD support the pro-angiogenic role of Nucleolin.

**Figure 7:**
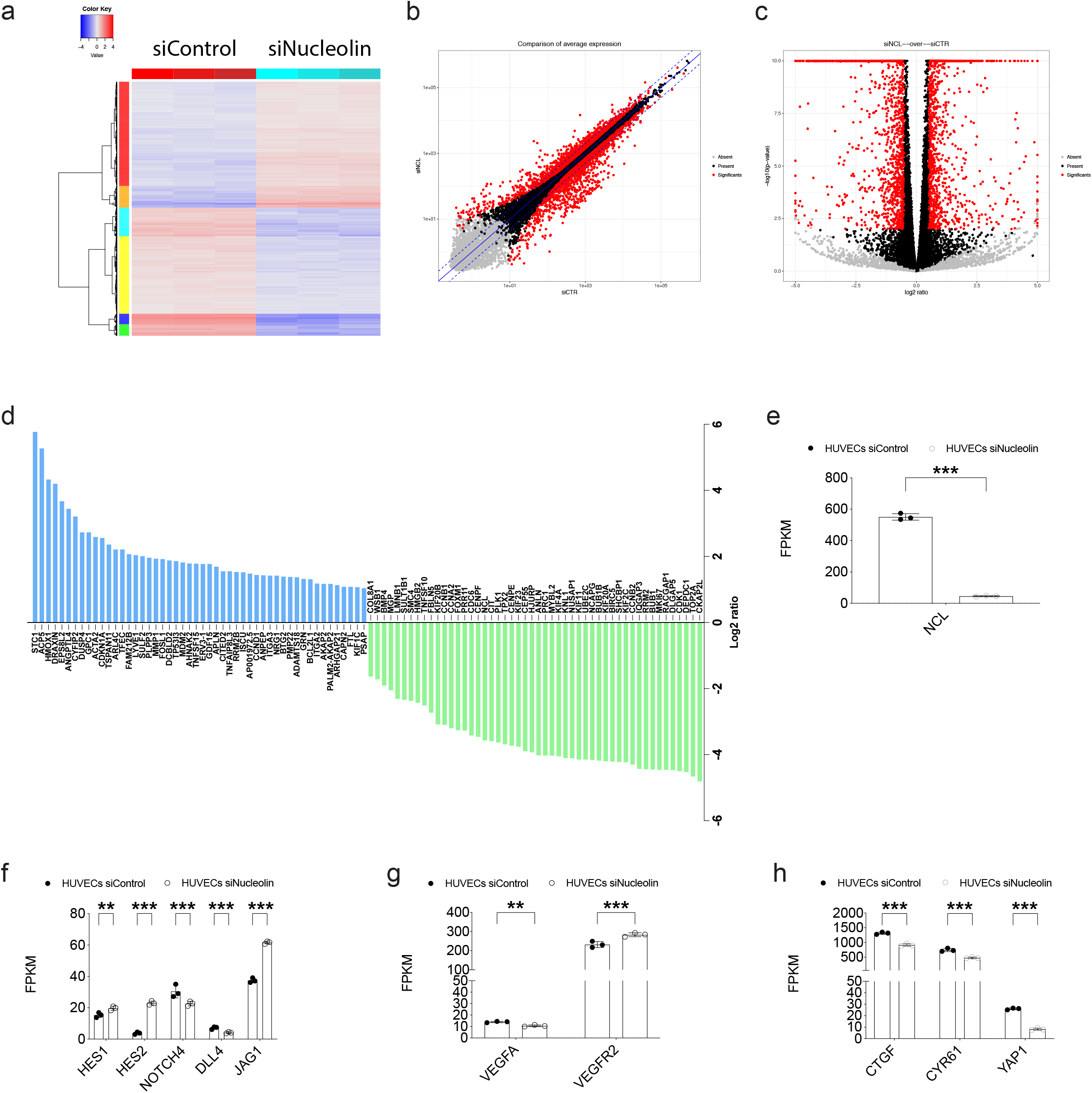
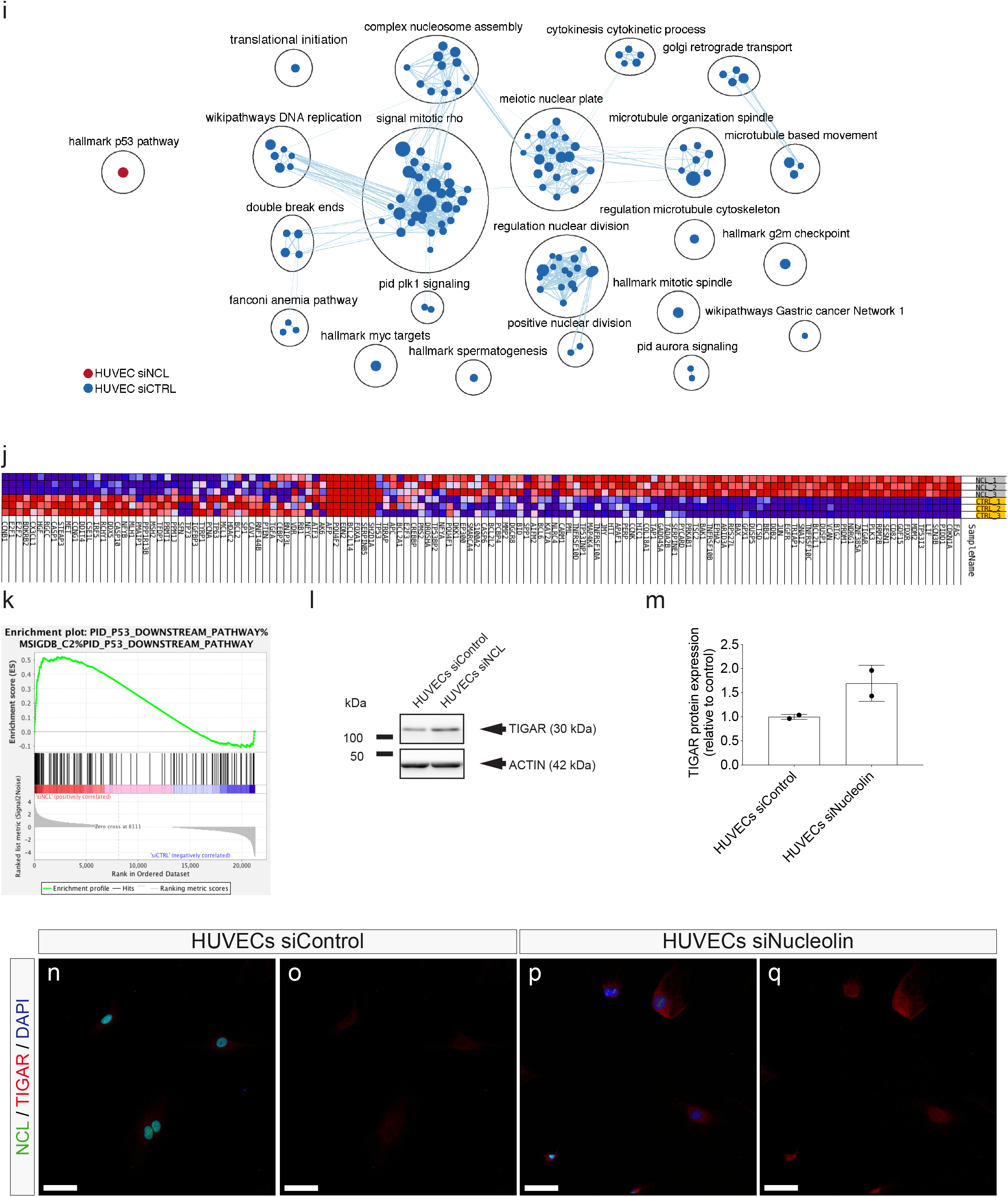
Nucleolin induces regulation of the Dll4-Jagged-Notch-Hey-Hes-NRARP-, the YAP-TAZ-CTGF-Cyr61- and the glucose (Hexokinase 2 (HK2), Glut1)- metabolic pathways through/via a P53-P21-TP53-inducible glycolysis and apoptosis regulator (TIGAR) pathway in HUVECs. Transcriptome analysis via RNA sequencing of HUVECs treated with small interfering RNA (siRNA) against Nucleolin (siNucleolin) and control siRNA (siControl) in three independent experiments. **a** Heat map and hierarchical clustering of siNucleolin treated HUVECs as compared to siControl treated HUVECs. **b-c** 3240 genes were differentially regulated between HUVECs^Nucleolin KD^ and HUVECs^Control KD^, and indicated in red on scatter (**b**) and volcano plots (**c**). **d** Top 50 significantly up (blue)- or down (green)-regulated genes detected by RNA-seq in HUVECs upon Nucleolin knock down as compared to control treatment. Differentially regulated genes were arranged according to fold change of gene expression. **e-h** Gene expression values (FPKM) of genes of interest for genes . Nucleolin was signigicantly downregulated by siNucleolin treatment (**e**). Nucleolin knock down induced a significant upregulation of the Dll4-Jagged-Notch signaling pathway including: Hes1, Hes2, Jagged1 (JAG1, **f**). NOTCH4, and DLL4 were downregulated upon Nucleolin knock down (**f**). VEGFA expression was downregulated and VEGFR2 expression upregulated (**g**) in HUVECs^Nucleolin KD^ as compared to HUVECs^Control KD^. siNucleolin treatment caused a significant down-regulation of the YAP-TAZ gene YAP1 as well as the YAP-TAZ downstream effectors genes CTGF and Cyr61 (**h**). **i-l** Gene set enrichment analysis (GSEA) and cytoscape enrichment map (**i**) showed a significant up-regulation of P53 signalling pathway in HUVECS treated with siRNA against Nucelolin. Whereas pathways involved in cell proliferation such as cell cycle regulation and mitosis were enriched in the control treatment. Pathways enriched in Nucleolin KD HUVECs are labeled in red and pathways enriched in control HUVECs are labeled in blue. Pathways are indicated by colored nodes. Their size represents the number of genes they contain. Green lines indicate relationships between the pathways. Black circles group related pathways. TIGAR was among the top 20 upregulated genes in the heat map grouping genes involved in P53 pathway (**j**). Enrichment plot indicates that P53 downstream pathways is enriched in Nucleolin KD HUVECs as compared to the control (**k**). siNucleolin treatment induced upregulation of P53 target genes TP53IP (**q**) and TIGAR (**r**) without affecting P53 expression (**l**). **m-n** Western blot showing TIGAR upregulation in HUVECs^Nucleolin KD^. Quantification of Western blot revealed an upregulation of TIGAR protein expression by almost 50% HUVECs^Nucleolin KD^. **o-r** HUVECs were stained for Nucleolin (green), TIGAR (red), and DAPI (blue). TIGAR expression was increased upon Nucleolin knockdown in HUVECs (**q**, **r**). \**P* < 0.05, \*\**P* < 0.01, \*\*\**P* < 0.001, \*\*\*\**P* < 0.0001.

Next, to further address the molecular pathways that are regulated by Nucleolin in endothelial cells, we performed a gene set enrichment analysis (GSEA)^83^ between siNCL-treatead HUVECs and the control HUVECs. Enrichment map revealed the TP53 pathway to be the only pathway enriched in Nucleolin knockdown HUVECs, whereas pathways downregulated in HUVEC^Nucleolin KD^ mainly belonged to cell cycle and proliferation (Fig. 7i), indicating the negative regulatory effect of Nucleolin on TP53 signaling and its positive regulatory effects on cell cycle and proliferation. Given the positive regulatory effects of Nucleolin on endothelial glucose metabolism and on cell cycle, we next wondered whether the expression of TP53-induced glycolysis and apoptosis regulator (TIGAR), a protein linking cell cycle and glucose metabolism^84,85^, was regulated upon Nucleolin-knockdown. Indeed, TIGAR was among the top 20 genes upregulated in the heatmap regrouping genes of the TP53 pathway (Fig. 7u). Knockdown of Nucleolin induced an upregulation of TP53 downstream signaling without affecting TP53 expression (Fig. 7). Namely, the expression of both TIGAR and the TP53 inducible protein 3 (TP53I3) were upregulated upon Nucleolin knockdown. Validation of these findings by Western blot and immunofluorescence confirmed Nucleolin-induced upregulation of TIGAR at the protein level (Fig. 7w-z_ii_).

Taken together, these results suggest that Nucleolin’s positive regulatory effects on CNS sprouting angiogenesis and endothelial metabolism might be regulated by a TP53-TIGAR-HK2 signaling axis.

## DISCUSSION

Here, using *in vitro* and *in vivo* approaches, we show that Nucleolin is a positive regulator of angiogenesis in the human fetal brain. Our results suggest that Nucleolin promotes endothelial sprouting, -proliferation, -filopodia formation, -and glucose metabolism via the TP53-TIGAR-HK2 pathway. We propose that, by acting on the cytoskeleton of CNS endothelial (tip- and stalk) cells and their filopodia, and by regulating vascular endothelial metabolism, Nucleolin controls the sprouting and filopodia extension of growing CNS blood vessels during human fetal brain development, and presumably in human brain tumors.

### Nucleolin - a putative NVU/PVN derived signal to regulate developmental brain- and brain tumor growth?

After initial development of the perineural vascular plexus (future meninges) via vasculogenesis, the CNS parenchyma is predominantly vascularized by sprouting angiogenesis ^58,86^. Most of the evidence regarding the molecular regulation of sprouting angiogenesis during brain development is based on murine studies, whereas even less knowledge exists how the vascularization and endothelial tip cells are regulated in the human brain.

Interestingly, angiogenesis is highly dynamic during brain development and almost quiescent in the adult healthy brain^15,18,59^, but is reactivated in a variety of angiogenesis-dependent CNS pathologies such as brain tumors, brain vascular malformations, or stroke^8,9,18^, thereby activating endothelial- and perivascular cells of the NVU^11,15,59^. In our study, we not only observed a reactivation of Nucleolin in angiogenic endothelial- and perivascular cells within glial brain tumors but also a positive correlation of Nucleolin expression with astrocytic tumor progression, suggesting a crucial role of Nucleolin as a perivascular niche-derived signal in both angiogenic- and tumor growth. The perivascular niche has been shown to activate tumor growth in mouse- and zebrafish models of breast cancer^28^. Notably, the stable microvasculature provided suppressive cues that inhibited angiogenesis, whereas the activated microvasculature fueled breast cancer cell growth via a direct cellular crosstalk between endothelial- and tumor cells. Strikingly, endothelial-derived thrombospondin-1 in the stable microvasculature induced sustained breast cancer cell quiescence but this suppressive cue was lost in sprouting neovasculature, where endothelial tip cell-derived active TGF-β1 and periostin promoted breast tumor growth^28^. These characterized the stable microvasculature as a “dormant perivascular (tumor) niche” in contrast to the sprouting neovasculature which constitutes an “activated perivascular (tumor) niche” in which endothelial tip cells exert crucial roles. In light of these studies, the exploration of Nucleolin angiogenic function within the perivascular tip cell niche *in vivo* promises to be an exciting avenue of future investigations.

Here, we found that Nucleolin is an important positive regulator of angiogenesis and endothelial tip cell filopodia in human fetal brain. The expression of Nucleolin in endothelial- and perivascular cells such as astrocytes and pericytes within the neurovascular unit of the human fetal brain, its downregulation in the adult healthy brain and reactivation in brain tumors suggests an integral role once reactivated during brain cancer. Its high expression in CD105^+^ angiogenic blood vessel endothelial cells in both human fetal brain and human glioblastoma (but not in the adult healthy brain) supports this presumed role in active (developmental, tumor) versus stable (adult healthy) brain angiogenesis. Accordingly, our *in vitro* results from a variety of functional bioassays suggest that by exerting stimulatory effects on sprouting angiogenesis, filopodia extension, and glucose metabolism of vascular endothelial cells, Nucleolin might promote the sprouting of angiogenic blood vessel endothelial cells into the brain parenchyma. The latter is also strongly suggested by the positive correlation between Nucleolin expression and the number of endothelial tip cell filopodia in the fetal brain parenchyma *in vivo*. Other neurodevelopmental regulators such as VEGF-A, and GPR124 are downregulated in the adult healthy CNS and are reactivated in vascular-dependent CNS pathologies such as brain tumors or stroke^87–89^. However, in contrast to these other pro- or anti-angiogenic factors, Nucleolin is - to the best of our knowledge - one of the first to have been directly described and compared in both human fetal brain and human gliomas. Furthermore, Nucleolin endothelial- and perivascular expression within the fetal- and tumor perivascular niche and its presumed different roles on the involved cell types (angiogenesis vs. tumor growth) characterizes Nucleolin as an important developmental signal that is reactivated during brain tumor growth.

### Expression patterns of Nucleolin in the developing and adult brain as well as in brain intrinsic tumors

Nucleolin is known to be expressed either at the cell surface, in the cytoplasm, nucleoplasm, or in the nucleolus^36,37,39^. At the cell surface, it functions mainly as a cell surface receptor for several ligands mediating various functions such as the growth of cancer cells or the apoptosis of endothelial cells^39,43,71,90^, whereas intracellularly, it accounts for transport of other proteins between the nucleus and the cytoplasm^36,39,90^.

Here, we observed that Nucleolin expression in the adult brain was mainly restricted to the nucleoli, as opposed to the predominant nucleoplasm expression during brain development and in brain tumors. It is well known that Nucleolin functions depend on its subcellular localization, which can be at the cell plasma membrane, in the cytoplasm, in the nucleus (nucleoplasm and/or nucleolus), and that nucleolar fraction of Nucleolin usually represents more than 90% of the cellular pool of Nucleolin^91^. The nucleolar restriction of Nucleolin in the adult brain may indicate that its main functions in the adult brain are those not linked to cell proliferation and cell division, such as ribosomal biogenesis and rRNA synthesis allowing for protein synthesis thereby supporting basic cellular functions^91 36,37,39^. However, during brain development and in brain tumors, perivascular- and endothelial cells are dividing at high rates and may thus require both, Nucleolin function in the ribosomes (in the nucleolus) as well as its regulatory role on cell cycle and cell proliferation (in the nucleoplasm) ^91^. Indeed, nuclear/nucleoplasmic Nucleolin is involved in the regulation of oncogene gene expression, promotes the proliferation and aggressiveness of a variety of tumors ^91^, and protects cancer cells from senescence ^91^.

In line with these observations, we observed a spatial restriction of Nucleolin to the nucleolus upon Nucleolin knocking-down *in vitro*, further indicating a restriction on specific basic functions in conditions with decreased Nucleolin expression (such as in the adult brain *in vivo*). Moreover, even though we observed nuclelar/nucleoplasmic and nucleolar expression, we did not observe cytosolic or cell surface/plasmalemmal expression of Nucleolin, which might be related to specific organ (e.g. brain versus periphery) and developmental (e.g. brain development, adult brain, brain tumors) properties. Given the complexity of Nucleolin’s subcellular expression patterns in various cell types^91^, the different functions of Nucleolin in the context of its cellular expression and specific subcellular localization await further investigations.

### Nucleolin as a regulator of endothelial filopodia dynamics and actin cytoskeleton orientation

Our *in vitro* results indicate that the effects of Nucleolin on developmental CNS angiogenesis are mediated via positive regulatory effects on endothelial sprouting, -proliferation, -glucose metabolism -and tip cell filopodia formation. This is supported by our *in vivo* findings showing Nucleolin expression in endothelial tip- stalk- and phalanx cells. Whereas our data indicate a role for Nucleolin in both tip cell filopodia formation as well as in stalk cell proliferation, the relative importance of Nucleolin for those angiogenic cell types and cellular mechanisms *in vivo* and *in vitro* remains unclear. For instance, differential expression patterns of Nucleolin might influence the competitiveness for the tip cell position^27,79^ and Nucleolin’s role on tip versus stalk cell specification could be tested using *in vitro* genetic mosaic sprouting assays^27^ and *in silico* computational simulation^79,92^. These future experiments might also shed light on the molecular mechanism that are responsible for our interesting observation of a positive correlation of Nucleolin expression and the number of filopodia extension per endothelial tip cell during brain development *in vivo*. It will be exciting to further investigate Nucleolin role on migrating tip cells, proliferating stalk cells and tip vs. stalk cell specification *in vivo* in both health and disease.

During sprouting angiogenesis and vessel branching, actin cytoskeleton orientation is crucial for endothelial tip- and stalk cell filopodia- and lamellipodia formation, sprout formation and guidance, and cell migration^15,23,24,59,93,94^. For instance, actin fibers form filopodia and lamellipodia in endothelial tip cells during sprout migration and actin polarizes in the direction of migration ^15,24,59,94^. The knock-down of Nucleolin in HUVECs lead to disorientation of the actin cytoskeleton inducing a loss of cell polarization as indicated by a change in cell shape towards a less elongated phenotype. Recent evidence suggests that MST1–FOXO1-mediated endothelial tip cell polarity facilitates sprouting angiogenesis^95^. MST1-FOXO1 was shown to be essential for directional migration of tip cells towards hypoxic regions and endothelial-specific deletion of either MST1 or FOXO1 lead to loss of endothelial tip cell polarity resulting in impaired sprouting angiogenesis^95^. Indeed, in our bulk RNA sequencing data, we found a downregulation of FOXO1 upon Nucleolin knock-down, indicating a regulatory role for Nucleolin on MST1-FOXO1-mediated regulation of cell polarity. However, MST1 was upregulated in the Nucleolin-deprived HUVECs, which is contradictory as compared to the findings by Kim and colleagues mentioned above. Whether these differences are due to *in vivo* versus *in vitro* conditions or mouse vs. human cells warrants future investigation. Also, it remains to be determined whether Nucleolin might regulate endothelial polarization via interaction with other signaling pathways in brain development and brain tumors *in vivo*.

Cell polarization directly depends on actin cytoskeleton organization^96^ and Nucleolin affected actin cytoskeleton orientation in our study. However, the molecular mechanisms behind the lost of actin cytoskeleton orientation upon knocking-down of Nucleolin remain to be elucidated. They could either be mediated via direct effects on the nucleolus given the well-known functions it exerts on angiogenic mechanisms such as cell migration, -sprouting, and lamellipodia- and filopodia formation^75,97^. Another possibility could be that Nucleolin affects protein-protein interactions important for actin cytoskeleton regulation such as for instance actin-myosin interaction. Interestingly, it was shown that glycolytic enzymes such as PFKFB3 associate with actin in endothelial filopodia and lamellipodia to provide the high amounts of ATP required for actin-myosin contraction^24,77^. Therefore, the decreased endothelial glycolysis that we observed in Nucleolin-deprived HUVECs might be causally linked to the decreased forced applied by those HUVEC^Nucleolin KD^ filopodia, e.g. due to reduced actin-myosin contraction. Furthermore, in light of emerging evidence of organ-specific regulation of angiogenesis^15,59,98^, it is tempting to speculate about an organ- or even CNS-specific regulation of Nucleolin on angiogenesis and endothelial tip cell polarity.

### Nucleolin as regulator of glucose-, but not fatty acid endothelial metabolism – effects on endothelial tip- and stalk cells?

Endothelial metabolism has recently emerged to be a crucial regulator of sprouting angiogenesis during development and in tumors^76–78,81,99,100^. Moreover, it was suggested that endothelial tip cells mainly rely on glycolysis whereas endothelial stalk cells also use fatty acid metabolism to support proliferation^76–78,81,99,100^. Here, we found that Nucleolin positively regulates endothelial glycolysis-but not fatty acid metabolism *in vitro*, indicating that Nucleolin’s main effect might be on tip cells. On the one hand, these findings are supported by the regulatory effects of Nucleolin on the number of filopodia during fetal brain development. In addition, the regulatory effect of Nucleolin on the actin cytoskeleton *in vitro* might – at least in part - explain the positive correlation between Nucleolin and the number of filopodia observed *in vivo*. Filopodia formation relies on actin cytoskeleton orientation and glycolytic production of ATP has been shown to promote filopodia formation *in vivo, in vitro*, and *in silico*^79^. On the other hand, given the observed *in vivo* expression of Nucleolin in both endothelial tip- and stalk cells as well as the positive regulatory effect on both sprouting angiogenesis (a tip cell function) and endothelial proliferation (a stalk cell function) *in vitro*, Nucleolin’s precise roles on both tip- and stalk cells in- and outside the CNS, for instance in the embryonic or postnatal brain^20,59^ or in the postnatal retina^101^ need to be investigated *in vivo*.

As demonstrated by our bulk RNA sequencing, most pathways regulated downstream of Nucleolin were linked to regulating cell cycle and cell proliferation, in agreement with Nucleolin’s well known role in these biological processes^37,43,90^. Interestingly, we also found a putative link between Nucleolin and endothelial metabolism via regulation of the TP53-TIGAR pathway. Previously, TIGAR was shown to regulate HK2 activity in tumor cells *in vitro*^102^ but an interaction of TIGAR with endothelial metabolism was not described so far. Our data suggest that Nucleolin downregulation promotes TP53-TIGAR signaling thereby inhibiting endothelial metabolism.

Moreover, the knock-down of Nucleolin lead to a downregulation of VEGF-A. The seemingly contradictory finding that VEGFR2 was upregulated in Nucleolin KD HUVECs may be due to the complexity of VEGF – VEGFR interactions also involving compensatory mechanism^103^ . In light of the important role of this signaling system for CNS angiogenesis ^58^ and endothelial tip cells ^8,18,104,105^, this result is interesting. It was showed that VEGF promotes Nucleolin translocation to the membrane *in vitro*^55^. However, whether and where the VEGF- and the Nucleolin signaling pathways interact intracellularly *in vivo* awaits further investigation. Moreover, given the crucial role of endothelial metabolism for vessel sprouting in development and disease^76,77,99^ as well as of tumor metabolism in gliomas^34^ investigating endothelial cell metabolism in brain tumors promises to be exciting.

### A putative role for Nucleolin in angiogenesis-dependent CNS pathologies?

Angiogenesis and the perivascular niche exert crucial roles in the pathophysiology of various vascular-dependent CNS diseases such as brain tumors, vascular malformations, and stroke ^11^ ^10,19,61^. Based on our findings, one may therefore speculate that Nucleolin could - in addition to brain tumors - also regulate vessel sprouting for example in brain vascular malformations such as arteriovenous- or cavernous malformations, or in neuroregenerative (e.g. stroke), or degenerative vascular diseases (e.g. vascular- or Alzheimer’s dementia). With regard to brain tumors, glycosylated surface Nucleolin has been been shown to increase with the malignancy grade of human gliomas^51^. A high expression of Nucleolin may therefore promote vascularization of astrocytoma and thereby promote brain tumor growth. Moreover, it was reported that delivery of a toxin directed to cell surface Nucleolin could be used as targeted therapy of human glioblastoma^106^. Another study reported that a Nucleolin antagonist induced death of primary human glioblastoma cells and decreased *in vivo* tumor growth in an orthotopic brain tumor model, but no regulatory effect on angiogenesis was addressed^107^. Furthermore the anticancer aptamer AS1411 - a G-rich quadruplex-forming oligodeoxynucleotide that binds specifically to Nucleolin - has shown promising clinical activity and is being widely used as a tumor-targeting agent^108^. Antibody- and peptide-mediated targeting of Nucleolin induced normalization of tumor vasculature in pancreas- and breast cancer models^56,72^, suggest strategies to explore effects of regulating Nucleolin on development of glioblastoma vasculature. These literature indications - in concert with our data - suggest that - in addition to its effect on tumor cell proliferation - Nucleolin may regulate brain tumor vascularization.

## ACKNOWLEDGMENTS

We thank our colleagues Giovanna Longo for help with fetal brain sample preparation for immunofluorescence and confocal imaging and Jiazhuo He for help with *in vitro* spheroid assays. The author(s) disclosed receipt of the following financial support for the research, authorship, and/or publication of this article: T.W. was supported by the OPO Foundation, the Swiss Cancer Research foundation (KFS-3880-02-2016-R, KFS-4758-02-2019-R), the Stiftung zur Krebsbekämpfung, the Kurt und Senta Herrmann Foundation, Forschungskredit of the University of Zurich, the Zurich Cancer League, the Theodor und Ida Herzog Egli Foundation, the Novartis Foundation for Medical-Biological Research, and the HOPE Foundation. K.D.B. was supported by grants of the Swiss National Science Foundation (31003A-176056) and a starting grant of the European Research council (716140). V.V. was supported by the Swiss National Science Foundation (SNF 310030B_133122/1), SNF NCCR “Molecular Systems Engineering”, the Commission of the European Communities/European Research Council (ERC) Advanced Grant (231157), FIRST and SCOPEM facility from ETH. D.V. was supported by Intramural Funding (2019).

## AUTHOR CONTRIBUTIONS

T.W. had the idea for the study, designed the experiments, wrote the manuscript, and made the figures. T.W., M.S., I.D.T., J-.Y.S., M.G., O.S., F.G., M.E., and M.Y. conducted the experiments and analyzed the data. R.S., V.V., D.V., K.D.B., and K.F. supervised the experiments in their respective labs, and T.W. supervised all the research. T.W. wrote the manuscript, M.S. helped editing the manuscript and figures. J.H., O.S., A.M.,, S.L., J.G., P.M., I.R., K.F., R.S., V.V., D.V., and K.D.B., gave critical inputs to the manuscript. All authors read and approved the final manuscript.

## MATERIALS AND METHODS

### Human fetal and adult brain tissue

Samples of fetal brain were obtained from three post-mortem fetuses, two 22- and one 18-week-old, derived from spontaneous abortions and received by the Department of Pathological Anatomy, University of Bari School of Medicine. Permission to collect fetal tissue was obtained from the mother at the end of the abortion procedure. The sampling and handling of the specimens conformed to the ethical rules of the Department of Emergency and Organ Transplantation, Division of Pathology, University of Bari School of Medicine, and approval was gained from the local Ethics Committee of the National Health System in compliance with the principles stated in the Declaration of Helsinki. The fetuses did not reveal macroscopic structural abnormalities at autopsy and/or microscopic malformations of the central nervous system after conventional histological analysis with H&E or toluidine blue staining. The fetal age was estimated based on the crown-rump length and/or pregnancy records (counting from the last menstrual period). From each fetus, samples of the dorso-lateral wall of the telencephalic vesicles (n=6; future cerebral hemispheres) were dissected along the coronal plane in slices about 0.5-cm thick, fixed for 2–3 hours at 4°C by immersion in 2% paraformaldehyde (PFA) plus 0.2% glutaraldehyde in phosphate-buffered saline solution (PBS, pH 7.6), washed in PBS and stored in PBS plus 0.02% PFA at 4°C. The parahippocampal cortex, used as normal adult brain samples obtained after selective amygdalohippocampectomy from patients with chronic pharmaco-resistant mesial temporal lobe epilepsy and glioblastoma samples were also cut in 0.5-cm thick slices and submitted to the same histological procedure applied to fetal slices.

### Immunofluorescence staining and analysis of angiogenesis and perivascular niche in human fetal brain, human adult brain, and human brain tumors

Fixed, unembedded fetal brain, normal adult brain, and glioblastoma slices were cut in 20-μm thick sections, using a vibrating microtome (Leica Microsystem) and submitted, as free-floating sections, for single and double staining, with the following antibodies (Abs): mouse mAb anti-nucleolin (1:200, Santa Cruz), rabbit pAb anti-CD31 (1:80; Abcam,), rabbit pAb anti-CD105 (predilute,; Abcam), rabbit pAb anti-GFAP (prediluted, Immunostar), pAb anti-NG2 (1:200, generous gift from William B. Stallcup). Briefly, the sections were: 1) permeabilized with 0.5% Triton X-100 in PBS for 30 min at room temperature (RT); 2) incubated overnight at 4°C with primary Abs, nucleolin, nucleolin/CD31, nucleolin/CD105, nucleolin/GFAP, and nucleolin/NG2; 3) incubated with the appropriate secondary Abs, goat anti-rabbit Alexa 568 and goat anti-mouse Alexa 488 (1:300, ThermoFisher Scientific,), for 45 min at RT; 4) counterstained with the nuclear dye TO-PRO-3 (diluted 1:10’000; Life Technologies, Inc., Gaithersburg, MD, USA). Finally, the sections were collected on polylysine slides (Menzel-Glaser, GmbH, Braunschweig, Germany) and coverslipped with Vectashield (Vector Laboratories Inc.). Negative controls were prepared by omitting the primary antibodies and mismatching the secondary antibodies. Sections were examined under Leica TCS SP5 confocal laser scanning microscope (Leica Microsystems) using a sequential scan procedure. Confocal images were taken at 0.35 μm intervals through the z-axis of the section, with 40x and 63x oil immersion lenses. Z-stacks of optical planes (image projection) and single optical planes were recorded and analysed by Leica confocal software

### Laser confocal morphometry

The quantitative assessment was carried out on fetal brains (n=3), normal brains (n=3) and glioblastomas (n=4) samples, by computer-aided morphometric analysis using the Leica Confocal Multicolor Package (Leica Microsystems) and ImageJ (NIH) softwares. The nucleolin mean area percentage was evaluated according to the ‘mean area fraction’ parameter of ImageJ, calculated on randomly chosen fields (n=10; field area = 150.000 μm^2^) of single optical planes across the z-stack. On double stained nucleolin/CD105 sections (fetal brain n=18, normal brain n=12, glioblastoma n=36), cells positive for both the markers were counted on maximum intensity projection images (40x magnification), cells positive for both nucleolin and CD31 were counted on double stained sections (fetal brain n=18, normal brain n=18, glioblastoma n=24), nucleolin^+^/GFAP^+^ cells were counted on double stained sections (fetal brain n=12, normal brain n=6, glioblastoma n=10), and nucleolin^+^/NG2^+^ cells were counted on double stained sections (fetal brain n=15, normal brain n=9, glioblastoma n=29). The percentage of double positive cells was calculated as the fraction of these cells on total number of nuclei stained with TO-PRO-3 counted on optical planes at 0.35μm intervals across the z-stack. The measurement of the number of vessel sprouts was carried out on nucleolin/CD31 and nucleolin/CD105 stained sections (n=3 for each fetus), to make the presence of endothelial tip cells recognizable. The results showed a range of 2-6 sprouts per section (total sprout number, n=35). The nucleolin green signal intensity was calculated on a manual selection of the cell nucleus as a ROI and then the number of filopodia emerging from each vessel sprout was recognized and counted on single optical planes acquired at 63x.

### Tissue Microarrays (TMAs)

Glioma tissue microarray. The study cohort comprised of low and high grade glioma samples from 103 patients who were treated at the Department of Neurosurgery, University Hospital Zurich (Switzerland) between 06/2003 and 05/2009. Paraffin blocks of these tumors were reviewed by a neuropathologist and classified according to the World Health Organization of brain tumors^66,67^. Representative tumor areas were marked on hematoxylin/eosin-stained slides by an experienced neuropathologist (JH) and two cores (0.6 mm diameter) were punched from the donor block and transferred into the tissue microarray (TMA) recipient block. Additionally, the TMA contained eight normal brain samples and four cell lines (MCF-7, LN-18, LN-229 and HT29).

For immunohistochemistry (IHC) analysis freshly cut 3-μm thick sections of the TMA block were mounted on SuperFrost. slides (Menzel Glaser, Braunschweig, Germany). The mouse monoclonal nucleolin antibody C23 (1:20, Santa Cruz) was used and incubated overnight at 4°C. As a detection system the Simple Stain MAX PO (MULTI) anti-mouse (Nichirei Biosciences) was used. Finally, the slides were counterstained with Mayer’s hemalum solution prior to dehydration and coverslipping.

Percentage of nucleolin staining of tumor cells in all the samples were evaluated by two investigators and was performed in an entirely blinded fashion.

### Cell culture and proliferation assay

Glioma cell lines (LN-229 and LN-18 kindly provided by N. de Tribolet, Geneva, Switzerland) and the ex vivo GBM-1 cells (passage 3-7) established at the Department of Neurosurgery, University Hospital Zurich, as described by Rodak et al. (J. Neurosurg 102: 1055-68, 2005) were cultured in DMEM (Gibco, Thermo Fisher Scientific, Allschwil, Switzerland) containing 10% heat-inactivated FCS (PAA Laboratories, Thermo Fisher Scientific), gentamicin (10 μg/ml, Gibco), 100 mM sodium pyruvate (ICN Biomedicals, Aurora, Ohio), and 5 ml N-acetyl L-alanyl L-glutamine (Biochrom AG, Berlin, Germany). Proliferation assays were performed by adding 1 μCi/ml of ^3^H-thymidine for the final 7 hours of culture and before harvesting and counting on a liquid scintillation counter (Wallac 1450 Microbeta/Trilux, Shelton, Connecticut). HUVECs were cultured in endothelial basal medium (EBM-2) supplemented with endothelial growth factors EGM-2 SingleQuots (Lonza).

### siRNA-mediated knockdown of Nucleolin in HUVECs

Silencing of Nucleolin expression, was performed by transfecting HUVECs with siRNA against human Nucleolin (50nM, Santa Cruz), or a control siRNA (Santa cruz), as indicated by the manufacturer (50 nM). HUVECs were transfected with the indicated siRNA using Transfection reagent and Transfection medium (Santa Cruz).

### Immunoblotting

Cells were lysed in RIPA buffer and mechanical disruption through a 1ml insulin syringe (BD). Proteins were separated by 10% SDS-polyacrylamide gel electrophoresis (SDS-PAGE) at 150 V and transferred to a nitrocellulose membrane at 100 V for 1.5 hours. Unspecific binding sites were blocked with 5% milk in TBS-Tween 0.05% for 30 minutes. The membrane was probed with mouse anti-Nucleolin (NCL, 1:500, Santa Cruz), rabbit anti-Hexokinase-2 (HK2, 1:500, proteintech), rabbit anti-6-phosphofructo-2-kinase/fructose-2,6-biphosphatase 3 (PFKFB3, 1:1,000, abcam), mouse anti-p53 (1:1,000, Cell signalling), rabbit anti-CPT1a (1:500, abcam), and rabbit anti-TIGAR (1:200, Sigma) at 4°C overnight. A secondary donkey antibody directed against mouse, or rabbit (1:2,000, Dako) was applied for 1 hour. Bands were visualised by chemiluminescence using ECL (Amersham). Densitometric analysis was performed with ImageJ (NIH freeware). Data were normalised to actin, and values of control cells were set to 1.

### Quantitative real-time PCR

Total RNA was prepared using the RNeasy RNA isolation kit (Qiagen, Hilden, Germany) including a DNase treatment to digest residual genomic DNA. For reverse transcription, equal amounts of total RNA were transformed by oligo(dT) and M-MLV reverse transcriptase (Promega). Ten nanograms of cDNA were amplified in the Applied Biosystems 7500 Fast Real Time PCR System thermocycler (Thermo Fisher Scientific) with the polymerase ready mix (KAPA SYBR FAST; Sigma). Relative quantification was calculated using the comparative threshold cycle (ΔΔ^CT^) method. cDNA levels were normalized to *S18* (reference genes) and a control sample (calibrator set to 1) was used to calculate the relative values.

### Spheroid angiogenesis assay

A three-dimensional (3D) *in vitro* sprouting angiogenesis assay was performed as described previously^73,77^. Briefly, HUVECs were incubated overnight in hanging drops in EGM-2 medium containing 20% methylcellulose (Sigma) to form spheroids. Spheroids were then embedded in collagen gel and cultured for 24 hours (at 37°C, 5% CO2) to induce sprouting. Spheroids were fixed with 4% PFA at room temperature for 15 min and images of spheroids were captured with a Leica DMi1 (objectives: 20x and 40x. Analysis of the number of sprouts and the average sprout length was done using Image J.

### Nanopillar arrays

The polymer nanopillars platforms (photoresistant SU-8 nanopillars, Micro Chem) were fabricated using nanosphere lithography followed by a molding process, as previously described ^74,75^. The spring constant k of a representative nanopillars was measured by atomic force microscopy (AFM). From the resulting force curve, k was calculated resulting in a value of 79 nN um^−1^. To enable HUVECs culture on nanopillars, arrays were placed on petri dishes. HUVECs on nanopillars deflected the nanopillar tips (displacement x) and the resulting forces exerted by MVECs on nanopillars could be dedicated using the formula F=k*x.

The nanopillar platforms were fabricated exploiting nanosphere lithography followed by a molding process [1] and further described in previous study [2]. The spring constant of a representative SU-8 nanopillar was measured by Atomic Force Microscopy (AFM). For the AFM measurement, a SU-8 single nanopillar was prepared by frozen sectioning and horizontal fixation. An AFM tip (of known spring constant 62 nN/μm) was used to measure the force curve of the SU-8 nanopillar. This experimentally determined pillar spring constant (*k*_AFM_ = 79 nN/μm [2], as given by the slope of the force-displacement curve, was then used to calculate the horizontal traction forces.

### Printing Fn on naopillar substrates for cell culture

The flat PDMS stamps were plasma treated for 5 seconds before use. 50 μg/ml fibronectin labelled with Alexa dye 633 were added to the stamp and was incubated at room temperature for 15 minutes. Excess Fn solution was removed and the stamp was dried by compressed air. Stamps coated with Fn were applied on top of the nanopillar substrates, 50g weight was applied on top to ensure good contact. The stamps were removed from nanopillar substrates and non-coated area was blocked with Pluronic F-127 (P2443, Sigma) solution. Control HUVEC and HUVEC Nucleolin KD were cultured in the EGM-2 medium (Lonza), at 37°C with 5% CO_2_. Cells (7,500 cells/cm^2^) were let to adhere for 6 h seeded on a SU-8 nanopillar array printed with fibronectin.

### Image acquisition and analysis

To calculate the traction forces by which the cells displaced the nanopillars, the deflections of the nanopillar tips were imaged with confocal microscopy using a Leica confocal microscope SP5 with a 63x/1.43 oil immersion objective. During image acquisition, cells were kept at 37 °C and 10% CO2 condition. The following laser wavelengths were used to acquire both the images of nanopillar arrays and of the labeled cells: 405 nm (DAPI), 488 nm (to create a DIC image of nanopillar structures, eGFP and FITC) and 546 nm (DiI & TRITC). Each cell was imaged live for 30 to 60 min, with a scanning ratio of 1 min/frame. The displacements in xy direction of the nanopillar tips were quantified by comparison of two images taken at the planes of the pillar tops versus the bottom plane, respectively, using confocal microscopy (image resolution 50 nm per pixel). All nanopillar images were processed by Diatrack 3.03 (Powerful Particle Tracking, Semasopht), Fiji (plugin, template matching for drift collection) and the force calculated according to Hooke’s law: F = *k*\*x. The average background displacement of pillars was ~15 nm.

### Scanning electron microscopy

Cells on nanopillar arrays were imaged with higher magnification using a Scanning Electron Microscope (SEM, Zeiss ULTRA 55). After fixation, the samples were dehydrated by critical point drying using standard procedures. Briefly, this stepwise dehydration procedure started by adding 0.5% aqueous solution of polyethylenimine, followed by dehydrating the samples with a series of ethanol-water washes (25%, 50%, 75%, two 95%, and 100% ethanol) and finishing by drying the sample using critical point drying equipment and CO_2_. After dehydration process, the samples were coated with a 5 nm thick layer of gold using a sputter coater.

### Cell immunofluorescence

Cells were cultured on nanopillars or 8-well chamber slides (Sigma-Aldrich), cells were fixed with 4% PFA (P6148, Sigma-Aldrich) for 15 min at room temperature (RT). The cell membrane was then permeabilized with 0.1% Triton X-100 (X100, Sigma-Aldrich) in PBS (10 min) followed by 1% BSA/PBS (85040C, Sigma-Aldrich) blocking step (30 min, RT).

HUVECs were stained with with mouse anti-NCL (1:100, Santa Cruz), rabbit anti-HK2 (1:100, proteintech), rabbit anti-PFKFB3 (1:100, abcam, rabbit anti-CPT1a (1:500, abcam), and rabbit andti-TIGAR (1:200, Sigma) in 1% BSA/PBS at 4°C overnight. Cells were the incubated with secondary antibodies goat anti-rabbit Alexa 568 and goat anti-mouse Alexa 488 (1:500, Thermo Fisher Scientific), or TRITC-labeled phalloidin (1:100 Sigma) in 1% BSA/PBS for 1.5 hours at RT. Nuclei were counterstained with DAPI staining using (1:20,000, Thermo Fisher Scientific) in PBS for 5 min.

### Glycolytic Flux

HUVECs were incubated for 6 hours in EGM-2 containing 0.4 μCi/ml [5-3H]-D-glucose (PerkinElmer). Supernatant was transferred into glass vials containing perchloric acid and sealed with rubber stoppers. ^3^H_2_O was captured in hanging wells containing filter paper soaked with H_2_O over a period of 48 hours at 37°C^80^. Thereafter, the filter paper was transferred in scintillation cocktail for radioactivity measurement by liquid scintillation counting.

### Fatty acid oxidation

HUVECs were incubated in FBS-free EGM-2 medium supplemented with 50 fatty-acid free BSA, 50 μM carnitine, 100 μM unlabeled palmitic acid, and 2 μCi/ml [9,10-^3^H]-palmitic acid. Again, supernatant was transfered into glass vials and sealed with rubber stoppers. Radioactivity was measured as in the glycolytic flux assay.

### Glucose uptake and lactate production

Glucose and lactate concentrations in HUVECs^Nucleolin KD^ and HUVECs^Control KD^ supernatant as well as in basal growth medium were measured at Toronto General Hospital.

Glucose uptake was calculated by substracting glucose concentration in medium by the concentration in cell supernatant. Lactate production was calculated by substracting lactate concentration in medium by the concentration in cell supernatant.

### Proliferation assay

HUVECs were incubated for 6 hr in growth medium supplemented with 1 μCi/ml [^3^H]-thymidine. Cells were fixed with 100% ethanol for 15 min at 4°C. Thereafter, cells were precipitated with 10% trichloroacetic acid and lysed using 0.1N NaOH. [^3^H]-thymidine incorporation into DNA was measured by scintillation counting.

### Statistical analysis

Statistical significance was determined using unpaired two-tailed Student’s t-test (GraphPad Prism8). Differences were considered significant with a P value less than 0.05. Quantified data are presented as mean ± SEM.

## REFERENCES

1 Singh, S. K. et al. Identification of human brain tumour initiating cells. Nature 432, 396–401, doi:10.1038/nature03128 (2004).

2 Chen, J. et al. A restricted cell population propagates glioblastoma growth after chemotherapy. Nature 488, 522–526, doi:10.1038/nature11287 (2012).

3 Thakkar, J. P. et al. Epidemiologic and molecular prognostic review of glioblastoma. Cancer Epidemiol Biomarkers Prev 23, 1985–1996, doi:10.1158/1055-9965.EPI-14-0275 (2014).

4 Batchelor, T. T., Reardon, D. A., de Groot, J. F., Wick, W. & Weller, M. Antiangiogenic therapy for glioblastoma: current status and future prospects. Clin Cancer Res 20, 5612–5619, doi:10.1158/1078-0432.CCR-14-0834 (2014).

5 Hegi, M. E. et al. MGMT gene silencing and benefit from temozolomide in glioblastoma. N Engl J Med 352, 997–1003, doi:10.1056/NEJMoa043331 (2005).

6 Stupp, R. et al. Radiotherapy plus concomitant and adjuvant temozolomide for glioblastoma. N Engl J Med 352, 987–996, doi:10.1056/NEJMoa043330 (2005).

7 Gilbert, M. R. et al. Dose-dense temozolomide for newly diagnosed glioblastoma: a randomized phase III clinical trial. J Clin Oncol 31, 4085–4091, doi:10.1200/JCO.2013.49.6968 (2013).

8 Carmeliet, P. & Jain, R. K. Molecular mechanisms and clinical applications of angiogenesis. Nature 473, 298–307, doi:10.1038/nature10144 (2011).

9 Jain, R. K. & Carmeliet, P. SnapShot: Tumor angiogenesis. Cell 149, 1408–1408 e1401, doi:10.1016/j.cell.2012.05.025 (2012).

10 Jain, R. K. et al. Angiogenesis in brain tumours. Nat Rev Neurosci 8, 610–622, doi:10.1038/nrn2175 (2007).

11 Hjelmeland, A. B., Lathia, J. D., Sathornsumetee, S. & Rich, J. N. Twisted tango: brain tumor neurovascular interactions. Nat Neurosci 14, 1375–1381, doi:10.1038/nn.2955 (2011).

12 Jayson, G. C., Kerbel, R., Ellis, L. M. & Harris, A. L. Antiangiogenic therapy in oncology: current status and future directions. Lancet 388, 518–529, doi:10.1016/S0140-6736 (15)01088-0 (2016).

13 Batchelor, T. T. et al. AZD2171, a pan-VEGF receptor tyrosine kinase inhibitor, normalizes tumor vasculature and alleviates edema in glioblastoma patients. Cancer cell 11, 83–95, doi:10.1016/j.ccr.2006.11.021 (2007).

14 Gilbert, M. R. et al. A randomized trial of bevacizumab for newly diagnosed glioblastoma. N Engl J Med 370, 699–708, doi:10.1056/NEJMoa1308573 (2014).

15 Walchli, T. et al. Wiring the Vascular Network with Neural Cues: A CNS Perspective. Neuron 87, 271–296, doi:10.1016/j.neuron.2015.06.038 (2015).

16 Weis, S. M. & Cheresh, D. A. Tumor angiogenesis: molecular pathways and therapeutic targets. Nat Med 17, 1359–1370, doi:10.1038/nm.2537 (2011).

17 Sweeney, M. D., Sagare, A. P. & Zlokovic, B. V. Blood-brain barrier breakdown in Alzheimer disease and other neurodegenerative disorders. Nat Rev Neurol 14, 133–150, doi:10.1038/nrneurol.2017.188 (2018).

18 Potente, M., Gerhardt, H. & Carmeliet, P. Basic and therapeutic aspects of angiogenesis. Cell 146, 873–887, doi:10.1016/j.cell.2011.08.039 (2011).

19 Arvanitis, C. D., Ferraro, G. B. & Jain, R. K. The blood-brain barrier and blood-tumour barrier in brain tumours and metastases. Nat Rev Cancer 20, 26–41, doi:10.1038/s41568-019-0205-x (2020).

20 Fantin, A., Vieira, J. M., Plein, A., Maden, C. H. & Ruhrberg, C. The embryonic mouse hindbrain as a qualitative and quantitative model for studying the molecular and cellular mechanisms of angiogenesis. Nat Protoc 8, 418–429, doi:10.1038/nprot.2013.015 (2013).

21 Ruhrberg, C. & Bautch, V. L. Neurovascular development and links to disease. Cell Mol Life Sci 70, 1675–1684, doi:10.1007/s00018-013-1277-5 (2013).

22 Segarra, M., Aburto, M. R., Hefendehl, J. & Acker-Palmer, A. Neurovascular Interactions in the Nervous System. Annu Rev Cell Dev Biol 35, 615–635, doi:10.1146/annurev-cellbio-100818-125142 (2019).

23 Gerhardt, H. et al. VEGF guides angiogenic sprouting utilizing endothelial tip cell filopodia. J Cell Biol 161, 1163–1177, doi:10.1083/jcb.200302047 (2003).

24 De Smet, F., Segura, I., De Bock, K., Hohensinner, P. J. & Carmeliet, P. Mechanisms of vessel branching: filopodia on endothelial tip cells lead the way. Arterioscler Thromb Vasc Biol 29, 639–649, doi:10.1161/ATVBAHA.109.185165 (2009).

25 Phng, L. K. & Gerhardt, H. Angiogenesis: a team effort coordinated by notch. Dev Cell 16, 196–208, doi:10.1016/j.devcel.2009.01.015 (2009).

26 Hellstrom, M., Phng, L. K. & Gerhardt, H. VEGF and Notch signaling: the yin and yang of angiogenic sprouting. Cell Adh Migr 1, 133–136 (2007).

27 Jakobsson, L. et al. Endothelial cells dynamically compete for the tip cell position during angiogenic sprouting. Nat Cell Biol 12, 943–953, doi:10.1038/ncb2103 (2010).

28 Ghajar, C. M. et al. The perivascular niche regulates breast tumour dormancy. Nat Cell Biol 15, 807–817, doi:10.1038/ncb2767 (2013).

29 Carlson, P. et al. Targeting the perivascular niche sensitizes disseminated tumour cells to chemotherapy. Nat Cell Biol 21, 238–250, doi:10.1038/s41556-018-0267-0 (2019).

30 Mazzone, M. et al. Heterozygous deficiency of PHD2 restores tumor oxygenation and inhibits metastasis via endothelial normalization. Cell 136, 839–851, doi:10.1016/j.cell.2009.01.020 (2009).

31 Bergers, G. & Benjamin, L. E. Tumorigenesis and the angiogenic switch. Nat Rev Cancer 3, 401–410, doi:10.1038/nrc1093 (2003).

32 Ribatti, D. & Djonov, V. Intussusceptive microvascular growth in tumors. Cancer Lett 316, 126–131, doi:10.1016/j.canlet.2011.10.040 (2012).

33 Frentzas, S. et al. Vessel co-option mediates resistance to anti-angiogenic therapy in liver metastases. Nat Med 22, 1294–1302, doi:10.1038/nm.4197 (2016).

34 Bi, J. et al. Altered cellular metabolism in gliomas - an emerging landscape of actionable co-dependency targets. Nat Rev Cancer 20, 57–70, doi:10.1038/s41568-019-0226-5 (2020).

35 Tajrishi, M. M., Tuteja, R. & Tuteja, N. Nucleolin: The most abundant multifunctional phosphoprotein of nucleolus. Commun Integr Biol 4, 267–275, doi:10.4161/cib.4.3.14884 (2011).

36 Ginisty, H., Sicard, H., Roger, B. & Bouvet, P. Structure and functions of nucleolin. J Cell Sci 112 (Pt 6), 761–772 (1999).

37 Tuteja, R. & Tuteja, N. Nucleolin: a multifunctional major nucleolar phosphoprotein. Crit Rev Biochem Mol Biol 33, 407–436, doi:10.1080/10409239891204260 (1998).

38 Srivastava, M. & Pollard, H. B. Molecular dissection of nucleolin’s role in growth and cell proliferation: new insights. Faseb J 13, 1911–1922 (1999).

39 Mongelard, F. & Bouvet, P. Nucleolin: a multiFACeTed protein. Trends Cell Biol 17, 80–86, doi:10.1016/j.tcb.2006.11.010 (2007).

40 Wolfson, E., Goldenberg, M., Solomon, S., Frishberg, A. & Pinkas-Kramarski, R. Nucleolin-binding by ErbB2 enhances tumorigenicity of ErbB2-positive breast cancer. Oncotarget 7, 65320–65334, doi:10.18632/oncotarget.11323 (2016).

41 Wolfson, E., Solomon, S., Schmukler, E., Goldshmit, Y. & Pinkas-Kramarski, R. Nucleolin and ErbB2 inhibition reduces tumorigenicity of ErbB2-positive breast cancer. Cell Death Dis 9, 47, doi:10.1038/s41419-017-0067-7 (2018).

42 Huang, F. et al. Phosphorylation of nucleolin is indispensable to its involvement in the proliferation and migration of non-small cell lung cancer cells. Oncol Rep 41, 590–598, doi:10.3892/or.2018.6787 (2019).

43 Xu, Z. et al. Knocking down nucleolin expression in gliomas inhibits tumor growth and induces cell cycle arrest. J Neurooncol 108, 59–67, doi:10.1007/s11060-012-0827-2 (2012).

44 Ishimaru, D. et al. Mechanism of regulation of bcl-2 mRNA by nucleolin and A+U-rich element-binding factor 1 (AUF1). J Biol Chem 285, 27182–27191, doi:10.1074/jbc.M109.098830 (2010).

45 Otake, Y. et al. Overexpression of nucleolin in chronic lymphocytic leukemia cells induces stabilization of bcl2 mRNA. Blood 109, 3069–3075, doi:10.1182/blood-2006-08-043257 (2007).

46 Takagi, M., Absalon, M. J., McLure, K. G. & Kastan, M. B. Regulation of p53 translation and induction after DNA damage by ribosomal protein L26 and nucleolin. Cell 123, 49–63, doi:10.1016/j.cell.2005.07.034 (2005).

47 Goldshmit, Y., Trangle, S. S., Kloog, Y. & Pinkas-Kramarski, R. Interfering with the interaction between ErbB1, nucleolin and Ras as a potential treatment for glioblastoma. Oncotarget 5, 8602–8613, doi:10.18632/oncotarget.2343 (2014).

48 Benedetti, E. et al. Nucleolin antagonist triggers autophagic cell death in human glioblastoma primary cells and decreased in vivo tumor growth in orthotopic brain tumor model. Oncotarget 6, 42091–42104, doi:10.18632/oncotarget.5990 (2015).

49 Luo, Z. et al. Precise glioblastoma targeting by AS1411 aptamer-functionalized poly (l-gamma-glutamylglutamine)-paclitaxel nanoconjugates. J Colloid Interface Sci 490, 783–796, doi:10.1016/j.jcis.2016.12.004 (2017).

50 Balca-Silva, J. et al. Nucleolin is expressed in patient-derived samples and glioblastoma cells, enabling improved intracellular drug delivery and cytotoxicity. Exp Cell Res 370, 68–77, doi:10.1016/j.yexcr.2018.06.005 (2018).

51 Galzio, R. et al. Glycosilated nucleolin as marker for human gliomas. J Cell Biochem 113, 571–579, doi:10.1002/jcb.23381 (2012).

52 Christian, S. et al. Nucleolin expressed at the cell surface is a marker of endothelial cells in angiogenic blood vessels. J Cell Biol 163, 871–878, doi:10.1083/jcb.200304132 (2003).

53 Shi, H. et al. Nucleolin is a receptor that mediates antiangiogenic and antitumor activity of endostatin. Blood 110, 2899–2906, doi:10.1182/blood-2007-01-064428 (2007).

54 Kadomatsu, K. & Muramatsu, T. Midkine and pleiotrophin in neural development and cancer. Cancer Lett 204, 127–143, doi:10.1016/S0304-3835(03)00450-6 (2004).

55 Huang, Y. et al. The angiogenic function of nucleolin is mediated by vascular endothelial growth factor and nonmuscle myosin. Blood 107, 3564–3571, doi:10.1182/blood-2005-07-2961 (2006).

56 Gilles, M. E. et al. Nucleolin Targeting Impairs the Progression of Pancreatic Cancer and Promotes the Normalization of Tumor Vasculature. Cancer Res 76, 7181–7193, doi:10.1158/0008-5472.CAN-16-0300 (2016).

57 Xu, C. et al. Targeting surface nucleolin induces autophagy-dependent cell death in pancreatic cancer via AMPK activation. Oncogene 38, 1832–1844, doi:10.1038/s41388-018-0556-x (2019).

58 Mancuso, M. R., Kuhnert, F. & Kuo, C. J. Developmental angiogenesis of the central nervous system. Lymphat Res Biol 6, 173–180, doi:10.1089/lrb.2008.1014 (2008).

59 Walchli, T. et al. Quantitative assessment of angiogenesis, perfused blood vessels and endothelial tip cells in the postnatal mouse brain. Nat Protoc 10, 53–74, doi:10.1038/nprot.2015.002 (2015).

60 Suzuki, T., Fujikura, K., Higashiyama, T. & Takata, K. DNA staining for fluorescence and laser confocal microscopy. J Histochem Cytochem 45, 49–53, doi:10.1177/002215549704500107 (1997).

61 Storkebaum, E., Quaegebeur, A., Vikkula, M. & Carmeliet, P. Cerebrovascular disorders: molecular insights and therapeutic opportunities. Nat Neurosci 14, 1390–1397, doi:10.1038/nn.2947 (2011).

62 Daneman, R., Zhou, L., Kebede, A. A. & Barres, B. A. Pericytes are required for bloodbrain barrier integrity during embryogenesis. Nature 468, 562–566, doi:10.1038/nature09513 (2010).

63 Armulik, A. et al. Pericytes regulate the blood-brain barrier. Nature 468, 557–561, doi:10.1038/nature09522 (2010).

64 Roitbak, T., Li, L. & Cunningham, L. A. Neural stem/progenitor cells promote endothelial cell morphogenesis and protect endothelial cells against ischemia via HIF-1alpha-regulated VEGF signaling. J Cereb Blood Flow Metab 28, 1530–1542, doi:10.1038/jcbfm.2008.38 (2008).

65 Chou, C. H., Sinden, J. D., Couraud, P. O. & Modo, M. In vitro modeling of the neurovascular environment by coculturing adult human brain endothelial cells with human neural stem cells. PLoS One 9, e106346, doi:10.1371/journal.pone.0106346 (2014).

66 Louis, D. N. et al. The 2016 World Health Organization Classification of Tumors of the Central Nervous System: a summary. Acta Neuropathol 131, 803–820, doi:10.1007/s00401-016-1545-1 (2016).

67 Perry, A. & Wesseling, P. Histologic classification of gliomas. Handb Clin Neurol 134, 71-95, doi:10.1016/B978-0-12-802997-8.00005-0 (2016).

68 Gerdes, J. et al. Cell cycle analysis of a cell proliferation-associated human nuclear antigen defined by the monoclonal antibody Ki-67. J Immunol 133, 1710–1715 (1984).

69 Ishii, N. et al. Frequent co-alterations of TP53, p16/CDKN2A, p14ARF, PTEN tumor suppressor genes in human glioma cell lines. Brain Pathol 9, 469–479, doi:10.1111/j.1750-3639.1999.tb00536.x (1999).

70 Diserens, A. C. et al. Characterization of an established human malignant glioma cell line: LN-18. Acta Neuropathol 53, 21–28, doi:10.1007/bf00697180 (1981).

71 Destouches, D. et al. Suppression of tumor growth and angiogenesis by a specific antagonist of the cell-surface expressed nucleolin. PLoS One 3, e2518, doi:10.1371/journal.pone.0002518 (2008).

72 Fogal, V., Sugahara, K. N., Ruoslahti, E. & Christian, S. Cell surface nucleolin antagonist causes endothelial cell apoptosis and normalization of tumor vasculature. Angiogenesis 12, 91–100, doi:10.1007/s10456-009-9137-5 (2009).

73 Korff, T., Krauss, T. & Augustin, H. G. Three-dimensional spheroidal culture of cytotrophoblast cells mimics the phenotype and differentiation of cytotrophoblasts from normal and preeclamptic pregnancies. Exp Cell Res 297, 415–423, doi:10.1016/j.yexcr.2004.03.043 (2004).

74 Kuo, C. W. et al. Polymeric nanopillar arrays for cell traction force measurements. Electrophoresis 31, 3152–3158, doi:10.1002/elps.201000212 (2010).

75 Shiu, J. Y., Aires, L., Lin, Z. & Vogel, V. Nanopillar force measurements reveal actin-cap-mediated YAP mechanotransduction. Nat Cell Biol 20, 262–271, doi:10.1038/s41556-017-0030-y (2018).

76 Eelen, G., Cruys, B., Welti, J., De Bock, K. & Carmeliet, P. Control of vessel sprouting by genetic and metabolic determinants. Trends Endocrinol Metab 24, 589–596, doi:10.1016/j.tem.2013.08.006 (2013).

77 De Bock, K. et al. Role of PFKFB3-driven glycolysis in vessel sprouting. Cell 154, 651–663, doi:10.1016/j.cell.2013.06.037 (2013).

78 Rohlenova, K., Veys, K., Miranda-Santos, I., De Bock, K. & Carmeliet, P. Endothelial Cell Metabolism in Health and Disease. Trends Cell Biol 28, 224–236, doi:10.1016/j.tcb.2017.10.010 (2018).

79 Cruys, B. et al. Glycolytic regulation of cell rearrangement in angiogenesis. Nat Commun 7, 12240, doi:10.1038/ncomms12240 (2016).

80 Aragones, J. et al. Deficiency or inhibition of oxygen sensor Phd1 induces hypoxia tolerance by reprogramming basal metabolism. Nat Genet 40, 170–180, doi:10.1038/ng.2007.62 (2008).

81 Schoors, S. et al. Fatty acid carbon is essential for dNTP synthesis in endothelial cells. Nature 520, 192–197, doi:10.1038/nature14362 (2015).

82 Kalucka, J. et al. Quiescent Endothelial Cells Upregulate Fatty Acid beta-Oxidation for Vasculoprotection via Redox Homeostasis. Cell Metab 28, 881–894 e813, doi:10.1016/j.cmet.2018.07.016 (2018).

83 Reimand, J. et al. Pathway enrichment analysis and visualization of omics data using g:Profiler, GSEA, Cytoscape and EnrichmentMap. Nature protocols 14, 482–517, doi:10.1038/s41596-018-0103-9 (2019).

84 Bensaad, K. et al. TIGAR, a p53-inducible regulator of glycolysis and apoptosis. Cell 126, 107–120, doi:10.1016/j.cell.2006.05.036 (2006).

85 Ko, Y. H. et al. TP53-inducible Glycolysis and Apoptosis Regulator (TIGAR) Metabolically Reprograms Carcinoma and Stromal Cells in Breast Cancer. J Biol Chem 291, 26291–26303, doi:10.1074/jbc.M116.740209 (2016).

86 Risau, W. Mechanisms of angiogenesis. Nature 386, 671–674 (1997).

87 Kuhnert, F. et al. Essential regulation of CNS angiogenesis by the orphan G protein-coupled receptor GPR124. Science 330, 985–989, doi:10.1126/science.1196554 (2010).

88 Chang, J. et al. Gpr124 is essential for blood-brain barrier integrity in central nervous system disease. Nat Med 23, 450–460, doi:10.1038/nm.4309 (2017).

89 Plate, K. H., Breier, G., Weich, H. A. & Risau, W. Vascular endothelial growth factor is a potential tumour angiogenesis factor in human gliomas in vivo. Nature 359, 845–848, doi:10.1038/359845a0 (1992).

90 Chen, Z. & Xu, X. Roles of nucleolin. Focus on cancer and anti-cancer therapy. Saudi Med J 37, 1312–1318, doi:10.15537/smj.2016.12.15972 (2016).

91 Jia, W., Yao, Z., Zhao, J., Guan, Q. & Gao, L. New perspectives of physiological and pathological functions of nucleolin (NCL). Life Sci 186, 1–10, doi:10.1016/j.lfs.2017.07.025 (2017).

92 Boas, S. E. & Merks, R. M. Tip cell overtaking occurs as a side effect of sprouting in computational models of angiogenesis. BMC Syst Biol 9, 86, doi:10.1186/s12918-015-0230-7 (2015).

93 Cao, J. et al. Polarized actin and VE-cadherin dynamics regulate junctional remodelling and cell migration during sprouting angiogenesis. Nat Commun 8, 2210, doi:10.1038/s41467-017-02373-8 (2017).

94 Carmeliet, P., De Smet, F., Loges, S. & Mazzone, M. Branching morphogenesis and antiangiogenesis candidates: tip cells lead the way. Nat Rev Clin Oncol 6, 315–326, doi:10.1038/nrclinonc.2009.64 (2009).

95 Kim, Y. H. et al. A MST1-FOXO1 cascade establishes endothelial tip cell polarity and facilitates sprouting angiogenesis. Nat Commun 10, 838, doi:10.1038/s41467-019-08773-2 (2019).

96 Lizama, C. O. & Zovein, A. C. Polarizing pathways: balancing endothelial polarity, permeability, and lumen formation. Exp Cell Res 319, 1247–1254, doi:10.1016/j.yexcr.2013.03.028 (2013).

97 Xu, Z. P., Tsuji, T., Riordan, J. F. & Hu, G. F. The nuclear function of angiogenin in endothelial cells is related to rRNA production. Biochem Biophys Res Commun 294, 287–292, doi:10.1016/S0006-291X(02)00479-5 (2002).

98 Herbert, S. P. & Stainier, D. Y. Molecular control of endothelial cell behaviour during blood vessel morphogenesis. Nat Rev Mol Cell Biol 12, 551–564, doi:10.1038/nrm3176 (2011).

99 Cantelmo, A. R. et al. Inhibition of the Glycolytic Activator PFKFB3 in Endothelium Induces Tumor Vessel Normalization, Impairs Metastasis, and Improves Chemotherapy. Cancer cell 30, 968–985, doi:10.1016/j.ccell.2016.10.006 (2016).

100 Eelen, G., de Zeeuw, P., Simons, M. & Carmeliet, P. Endothelial cell metabolism in normal and diseased vasculature. Circ Res 116, 1231–1244, doi:10.1161/CIRCRESAHA.116.302855 (2015).

101 Pitulescu, M. E., Schmidt, I., Benedito, R. & Adams, R. H. Inducible gene targeting in the neonatal vasculature and analysis of retinal angiogenesis in mice. Nat Protoc 5, 1518–1534, doi:10.1038/nprot.2010.113 (2010).

102 Cheung, E. C., Ludwig, R. L. & Vousden, K. H. Mitochondrial localization of TIGAR under hypoxia stimulates HK2 and lowers ROS and cell death. Proc Natl Acad Sci U S A 109, 20491–20496, doi:10.1073/pnas.1206530109 (2012).

103 Apte, R. S., Chen, D. S. & Ferrara, N. VEGF in Signaling and Disease: Beyond Discovery and Development. Cell 176, 1248–1264, doi:10.1016/j.cell.2019.01.021 (2019).

104 Blanco, R. & Gerhardt, H. VEGF and notch in tip and stalk cell selection. Cold Spring Harb Perspect Med 3, doi:10.1101/cshperspect.a006569 (2013).

105 Geudens, I. & Gerhardt, H. Coordinating cell behaviour during blood vessel formation. Development 138, 4569–4583, doi:10.1242/dev.062323 (2011).

106 Dhez, A. C. et al. Targeted therapy of human glioblastoma via delivery of a toxin through a peptide directed to cell surface nucleolin. J Cell Physiol 233, 4091–4105, doi:10.1002/jcp.26205 (2018).

107 Cheung, H. C. et al. Splicing factors PTBP1 and PTBP2 promote proliferation and migration of glioma cell lines. Brain 132, 2277–2288, doi:10.1093/brain/awp153 (2009).

108 Reyes-Reyes, E. M., Salipur, F. R., Shams, M., Forsthoefel, M. K. & Bates, P. J. Mechanistic studies of anticancer aptamer AS1411 reveal a novel role for nucleolin in regulating Rac1 activation. Mol Oncol 9, 1392–1405, doi:10.1016/j.molonc.2015.03.012 (2015).

